# Human Visual Search Follows Suboptimal Bayesian Strategy Revealed by a Spatiotemporal Computational Model

**DOI:** 10.1101/779652

**Authors:** Yunhui Zhou, Yuguo Yu

## Abstract

Humans perform sequences of eye movements to search for a target in complex environment, but the efficiency of human search strategy is still controversial. Previous studies showed that humans can optimally integrate information across fixations and determine the next fixation location. However, their models ignored the temporal control of eye movement, ignored the limited human memory capacity, and the model prediction did not agree with details of human eye movement metrics well. Here, we measured the temporal course of human visibility map and recorded the eye movements of human subjects performing a visual search task. We further built a continuous-time eye movement model which considered saccadic inaccuracy, saccadic bias, and memory constraints in the visual system. This model agreed with many spatial and temporal properties of human eye movements, and showed several similar statistical dependencies between successive eye movements. In addition, our model also predicted that the human saccade decision is shaped by a memory capacity of around 8 recent fixations. These results suggest that human visual search strategy is not strictly optimal in the sense of fully utilizing the visibility map, but instead tries to balance between search performance and the costs to perform the task.

**Author Summary:** During visual search, how do humans determine when and where to make eye movement is an important unsolved issue. Previous studies suggested that human can optimally use the visibility map to determine fixation locations, but we found that such model didn’t agree with details of human eye movement metrics because it ignored several realistic biological limitations of human brain functions, and couldn’t explain the temporal control of eye movements. Instead, we showed that considering the temporal course of visual processing and several constrains of the visual system could greatly improve the prediction on the spatiotemporal properties of human eye movement while only slightly affected the search performance in terms of median fixation numbers. Therefore, humans may not use the visibility map in a strictly optimal sense, but tried to balance between search performance and the costs to perform the task.

## Introduction

For animals with foveated retina, efficient visual search is important for survival, and requires close-to-optimal planning of eye movement both in space and time. Currently, there are conflict evidences on whether human can make spatially optimal eye movements. The Bayesian ideal searcher and the closely-related entropy-limit-minimization (ELM) model are two important optimal eye movement models of multi-saccade visual search task, and previous studies have shown that humans’ search performance and eye movement statistics were consistent with these optimal models [1–3]. However, other studies have indicated that human may rely more on suboptimal strategy rather than calculating and planning optimal eye movement for a specific task [4–7]. What’s more, despite the claim that human eye movement strategy was close to optimal, humans seemed to made fewer long saccades compared to the optimal models [1]. Humans also showed statistical dependencies between successive eye movement during visual search [8], but detailed comparison to the optimal models is still missing.

The Bayesian ideal searcher and the ELM model were optimal in the sense of fully using the visibility map [1, 3], but they didn’t consider other costs to perform the task. For humans, the requirement of unlimited memory capacity in these optimal models is unrealistic, and making eye movements also have costs. For example, longer saccades are less accurate and increase the chance to make a secondary saccade [9–13], take longer to finish [14–16], and disrupt vision during saccade more severely [17], so this might explain why humans made fewer long saccades than the optimal models [1]. With these constraints, it is unlikely that humans strictly follow the optimal search rule. We therefore reconsider the question of how optimal do humans search and ask if a visual search model with more biological limitations can perform closer to humans.

Moreover, the temporal control of eye movements has received surprisingly little attention in visual search models. For example, the Target Acquisition Model does not explain fixation duration [18, 19]; the Bayesian ideal searcher and ELM model assume each fixation last for 250 ms [1, 3]; and the Guided Search model assumes fixation duration is 200 – 250 ms [20]; whereas the fixation duration observed in experiment usually has a large variation ranging from less than 100 ms to more than 700 ms [21]. We found only one model of visual search that considered mean fixation duration, but it couldn’t simulate the large variation [22]. The lack of fixation durations in the above models may be caused by the view that visual search is more about deciding fixation locations in the environment to find the target than deciding how long each fixation should be. However, since interdependency between fixation duration and spatial control of eye movements has been observed [8,23,24], a complete model of visual search should explained control both in space and time instead of focusing on a single aspect.

Eye movements are typically viewed as sequence of decisions, and fixation duration can be viewed as reaction time of a saccade decision. Indeed, the distribution of fixation duration showed similar right-skewed shape as the reaction time distribution of other decision-making tasks [12,25,26]. Therefore, some models of reading and scene viewing capture the fixation duration distribution by using the drift-diffusion model of decision making [27–30], which is successful in explaining the reaction time distribution [31, 32]. The central idea is to accumulate information stochastically within a fixation and triggered a saccade whenever the accumulation reaches a threshold, which has receive wide experiment support [33–35]. Although this approach is successful, the accumulation process in these models are usually simplified or artificial as they did not quantitatively measure the dynamics of information accumulation within a fixation. Since previous researches have shown that visual search task difficulty affects fixation duration [36, 37], we believe that the information accumulation process should be explicitly measured according to the visual search stimulus used in the task, and the data can be incorporated to previous optimal models [1, 3] based on signal detection theory.

In this research, we experimentally measured the accumulation of visual information by the brief stimulus exposure method. The data was then used to formulate a drift-diffusion process that decides saccade timing. Combined with the ELM model, we proposed a continuous-time eye movement model that determines both fixation location and fixation duration. Though the search performance of human subjects was close to optimal, we found that their eye movement statistics was inconsistent of the optimal search strategy. Instead, adding constraints on saccade accuracy, saccade amplitude and memory capacity could significantly improve the model’s prediction on human eye movement statistics while only slightly impair the search performance.

## Results

### Selecting Target Contrast

The stimulus image was a Gabor target (6 cycle/degree, 0.3 degrees in diameter) embedded in a circular naturalistic background noise image (15 degrees in diameter, root-mean-squared (RMS) contrast 0.2) (Fig 1A). Ten student subjects were recruited in this study. To normalize visual search task difficulty across subjects, we first measured the relationship between foveal target visibility (*d’* in signal detection theory) and target RMS contrast. The subjects’ correct response rate could be well fitted by a Weibull psychometric function given the target RMS contrast (Fig 1B). Visibility was positively related to the target RMS contrast and could be described by Equation (2) (Fig 1C). For each participant, we selected the target RMS contrast that made visibility equals to 3.0 (Fig 1C). The selected value ranged from 0.111 to 0.135, with a mean value of 0.125.

**Fig 1.**
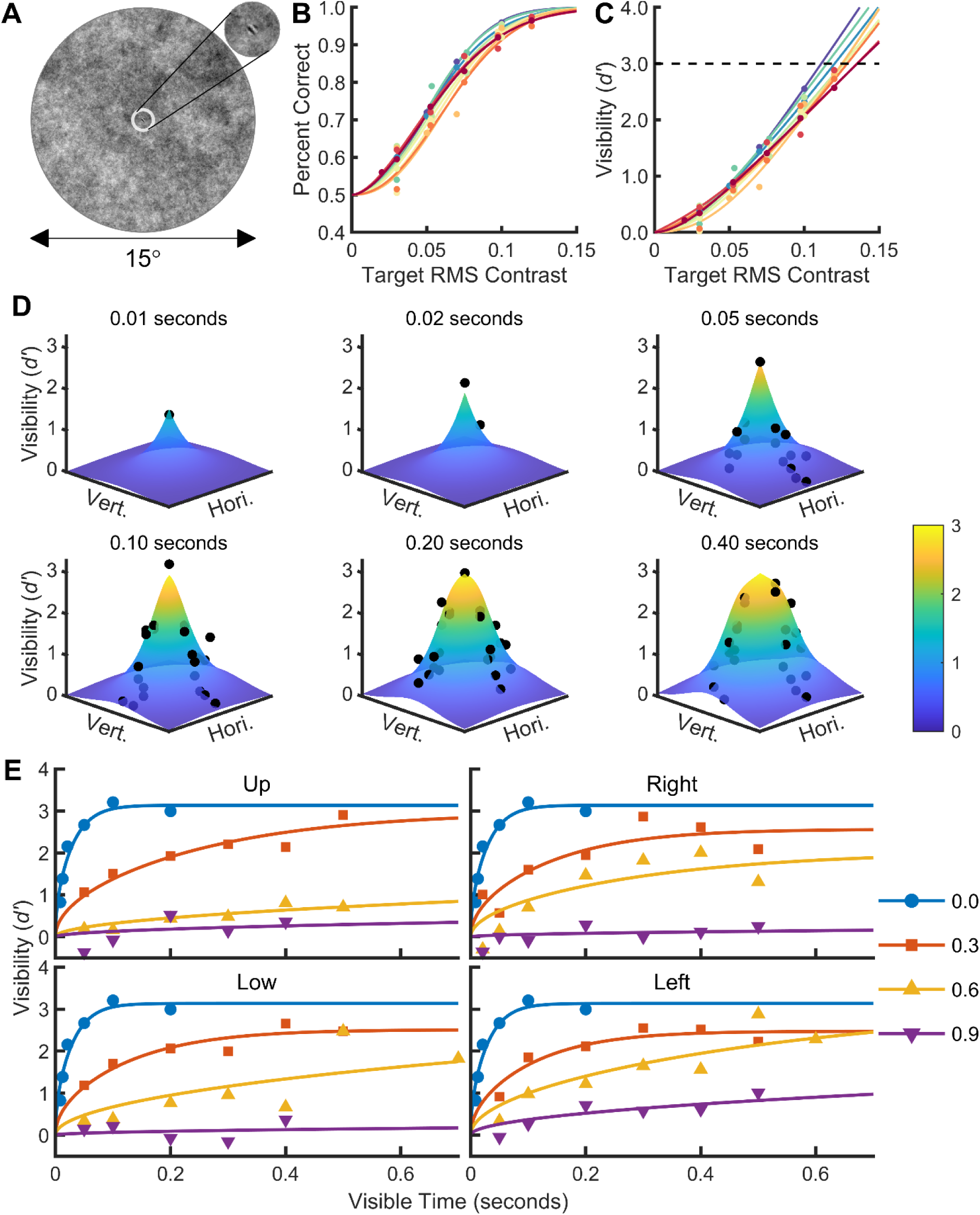
Stimulus image and results of the detection task. **A.** The stimulus image was a 1/f noise image (15° diameter, root-mean-squared (RMS) contrast 0.2) with an embedded target. The target was a 6 cycle/degree Gabor grating with a diameter of 0.3 degrees, oriented 45° counterclockwise from the vertical and windowed by a symmetrical raised cosine (half-height width of one cycle). The area outside the image was set to the mean gray value of the image (0.5, not shown here). **B, C.** Detection rate (B) and foveal target visibility (C) as a function of the target RMS contrast of the ten subjects. Dots are raw data (one color per participant). In B, the curves are the Weibull function fit to the dots. In C, the curves are calculated from Weibull functions fit to hit rates and the correct rejection rate of the raw data (Equation (1)). The intersection between the dashed line (d’ = 3.0) and the curves is the target RMS contrast chosen for each participant, which was used for all subsequent experiments. **D.** Target visibility map after different stimulus exposure times in the detection task. The dots are raw data merged from four subjects in first version of the detection task. The surface is the visibility map function (Equations (10), (12), (13)) fit to the raw data. Not all locations were tested in all exposure times so the number of black dots may differ in each subplot. Hori/Vert: horizontal/vertical dimension of the search field. Fixation location was at the center of search field. **E.** Temporal course of target visibility at four cardinal directions (shown in four subplots) relative to the fixation location. The dots are raw data merged from four subjects in first version of the detection task. Different markers represent raw data measured at different relative eccentricities (shown in the legend). The eccentricity values (0.0 - 0.9) are normalized with respect to the radius of the search field. The curves are Equation (10) fitted to the raw data measured at each location. The data measured at the fixation location (eccentricity = 0.0) are shown in every subplot. For clarity, not all measured locations are shown here. See Fig S1-S4 for the complete data from each participant.

### Temporal Course of Target Visibility and the Evidence Accumulation Model

We then measured how target visibility changed over time within one fixation across the visual field by a brief stimulus presentation experiment. Four of the ten subjects finished this experiment, and their data were pooled together. The target visibility first increased as a function of time, and then reached stable peak values after stimulus onset (Fig 1DE). The steady-state foveal visibility was approximately 3.14, which was close to the pre-defined value of 3.0. The steady-state and rising speed of visibility was higher in central visual field than in peripheral visual field (Fig 1E). There were some individual variability on the span of visibility map (e.g. some participant could detect the target at larger eccentricity), but the trend that central vision could detect the target faster than peripheral vision was basically preserved across participants (Fig S1 to Fig S4).

We then formulated an evidence accumulation model to incorporate the temporal course of visibility data into visual search models. We assumed that in each short time interval, the visual system received a small normally distributed random sample of visual information at each location. Similar to previous visual search models [1–3], the mean of the random variable depends on target’s presence at the location, but the variance is equal (Equation (3)). We further assumed that these random independent samples were integrated over time in a leaky manner (earlier samples were given lower weight, Equation (6)), because perfect integration could produce arbitrary high visibility given enough samples (Equation (4)). The integration of random samples produced a drift-diffusion process, and the distribution at each time point could be analytically described (Equations (7)-(9)). According to signal detection theory, visibility (*d’*) is the separation between the noise and target+noise distributions in units of their common standard deviation [38], therefore the temporal course of visibility could be described as Equations (10), (12), (13), and fit well to visibility data measured at each location (Fig 1DE).

### Subjects’ Eye Movement Statistics Were Predicted Better By A Suboptimal Visual Search Strategy

Next, we performed a visual search experiment with the same stimulus images to compare their eye movement data to visual search models. All ten subjects finished this task, and they were split into training set (the four subjects that measured the temporal course of visibility) and testing set (the rest six subjects).

We used the entropy-limit-minimization (ELM) model of visual search as the baseline model and it served as a replication of previous study [1]. To model eye movements control both in space and time, we combined the evidence accumulation model and the ELM model into a continuous-time ELM (CTELM) model. Since several previous studies have suggested that the control of fixation duration depends more on foveal than peripheral visual analysis [39, 40], the CTELM model terminates a fixation when the posterior probability of target being at current fixation location drops below a collapsing threshold (Fig 2). The next fixation location was chosen by the ELM rule (Equation (17)).

**Fig 2.**
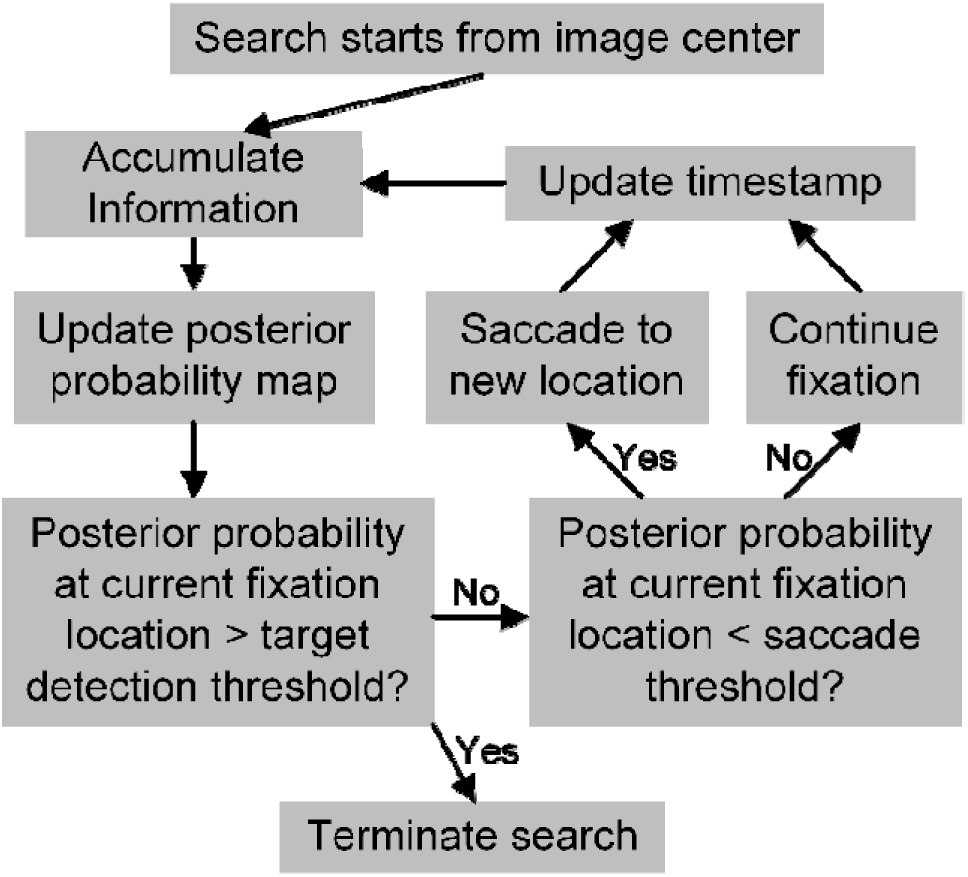
Schematic diagram of the continuous-time entropy limit minimization (CTELM) and constrained-CTELM (CCTELM) models.

We also tested whether adding constraints to the CTELM model (called the CCTELM model) could improve model’s prediction on experiment data. Three constraints were considered: preference for short saccades, inaccuracy of saccade landing position, and memory limitation. We applied a penalty function (Equation (25)) when calculating the expected information gain of the next fixation to suppress the probability of choosing a distant saccade target. The relationship between variability of saccade landing position and saccade target eccentricity was obtained from [12] (Fig S7), so that the model could fixate at any location instead of the predefined 400 locations (Fig S6). We also considered a simple form of memory so that the model could only integrate a fixed number of fixations when calculating the posterior probability of target location (Equation (23)). Visual information prior to what the memory could hold was discarded. We found that human eye movements predicted that the memory capacity was about 8 fixations (including the current fixation). In the next section we will show how we estimated the capacity.

There is only one free parameter in the ELM model that controls correct response rate, and three additional free parameters in the CTELM model that controls the fixation duration. The CCTELM model has five free parameters: one for correct response rate, three for fixation duration, and one for the saccade amplitude penalty function (see methods and Table S1). These parameters were fit to the training set and tested against the testing set.

We first examined how the three models predict subjects’ spatial control of eye movements. In previous studies of similar experiment, humans produced a doughnut-shaped fixation location distribution peaked at about 5° from image center, and fixated more at vertical part of the image than the horizontal part [1, 2]. Our subjects in testing set, however, fixated more uniformly across the image in both correct (Fig 3ACD) and error trials (Fig S8ACD), though similar to previous results more fixations were located in the vertical part (Fig 3B, Fig S8B). There was a fixation hotspot located slightly above the image center in experiment data (Fig 3A), but it was caused by the first saccade from two subjects, so we chose to focus more on global trends instead of this individual variability. All three models could predict the preference to fixate at vertical part of the image (Fig 3B, Fig S8B); however, the ELM and CTELM models produced a sharp doughnut-shaped distribution of fixation locations, whereas the CCTELM model fixated more uniformly and was similar to subjects’ data (Fig 3ACD, Fig S8ACD). Importantly, ELM and CTELM models progressively fixated at the outer part of the image as search continues, whereas the subjects’ and CCTELM model’s fixation distance to image center was relatively stable (Fig 3E, Fig S8E). Therefore, the agreement of ELM and CTELM model to experiment data decreased as the search progressed, but the CCTELM model could achieve stable and better agreement (Fig 3F, Fig S8F).

**Fig 3.**
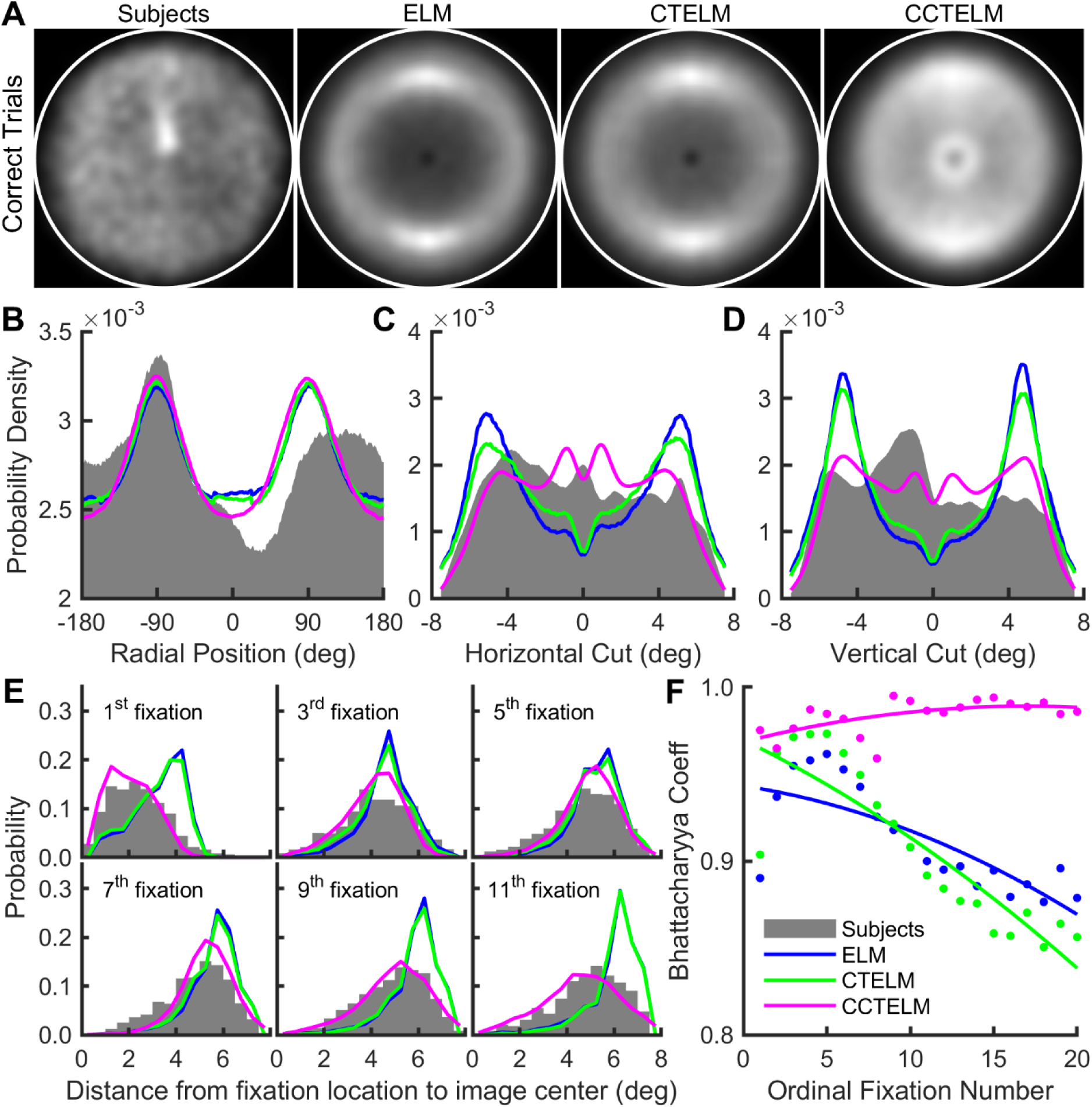
Fixation location distribution of subjects and models in correct trials (for error trials see Fig S8). **A.** Distribution of fixations location in the search field (inside the white circle), lighter means higher density. The densities were obtained by smoothing the scatterplot of the fixation locations by a Gaussian window with a standard deviation of 0.35 degrees (15 pixels) and then normalizing the maximum value in each subplot to one. **B.** Direction histograms of fixation location relative to the image center. The histograms were obtained with a sliding radial window with a width of 45° centered different radial positions (x-axis). Rightward direction is 0° and upward direction is −90°. **C.** Horizontal cuts (through the center) through the fixation densities in panel A. Rightward direction is positive in x-axis. **D.** Vertical cuts (through the center) through fixation densities in panel A. Upward direction is negative in x-axis. In panels BCD area under curve are normalized to one. **E.** Fixation distance distribution to image center selected within the initial 11 fixations after the first saccade. **F.** Bhattacharyya coefficient between models’ and subjects’ distributions of fixation distance to image center of the initial 20 fixations after the first saccade. Dots represent raw data; curves represent quadratic functions fitted to the dots. Legends from panel B to F are shown in bottom-left of panel F.

For the saccade amplitude, the two optimal models generated more long saccades than the subjects did in both correct and error trials (Fig 4A), which was similar to results in previous studies [1]. The difference was present since the first saccade (Fig S9, Fig S10), and the disparity between the models’ and humans’ distributions increased as the search progressed (Fig 4B). The CCTELM model prefer shorter saccades, and the prediction on experiment data was better and more stable (Fig 4AB). Besides, in a sequence of eye movements, the change of saccade direction after a saccade was usually larger for the ELM and CTELM models than CCTELM model and subjects (Fig 4C). Both subjects and the three models tended to make larger saccades after larger change in saccade direction, but the slope of the relationship was larger for both the ELM and CTELM than subjects and CCTELM model (Fig 4D). What’s more, both the ELM and CTELM models tended to make longer saccades after a long saccade, but the subjects and CCTELM model showed the opposite behavior (Fig 4E). Therefore, the CCTELM model could better explain the subjects’ spatial control of eye movements.

**Fig 4.**
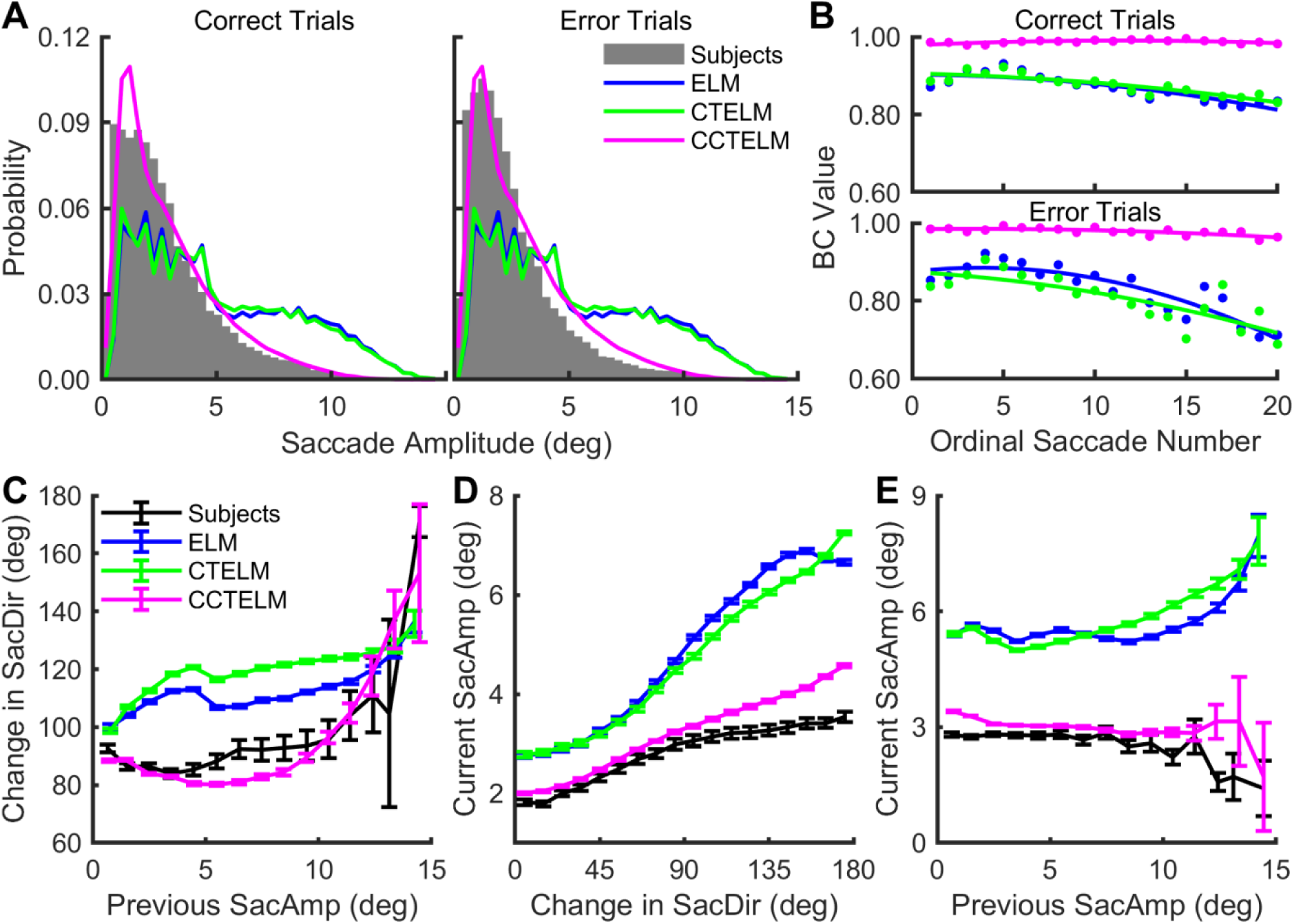
Saccade amplitude distribution of the subjects and models. **A.** Saccade amplitude histogram of all saccades in correct and error trials. **B.** Bhattacharyya coefficient (BC) between the models’ and subjects’ saccade amplitude distribution of the initial 20 saccades in correct and error trials. Dots represent raw data, curves represent quadratic functions fitted to the dots. See Fig S9 and Fig S10 for the distribution of raw data at each ordinal position in correct and error trials. **C.** Relationship between the first saccade’s amplitude (SacAmp) in two consecutive saccades and the change in saccade direction (SacDir) in all trials. **D.** Relationship between the second saccade’s amplitude in two consecutive saccades and the change in saccade direction in all trials. **E.** Relationship between the second and the first saccade’s amplitude in two consecutive saccades in all trials. In panels CDE error bar represents 95% confidence interval.

We next examined how CTELM and CCTELM models predict subjects’ temporal control of fixations. Both models could predict the overall fixation duration distribution (Fig 5A), and the agreement was stable as search progressed in both correct and error trials (Fig 5B, Fig S11, Fig S12). However, as the spatial and temporal control of eye movements are not independent, we then checked the relationship between fixation duration and other eye movement metrics. Humans tended to fixate longer when the directions of previous and next saccade were perpendicular to each other, forming an inverted-U shape relationship between fixation duration and change in saccade direction (Fig 5C). The CCTELM model could partially generate this relationship, but for the CTELM model, fixation tended to be longest when the directions of previous and next saccade form a 135° angle (Fig 5C). The reason for this behavior of the CCTELM model was because the previous saccade tended to be longer when the next saccade changed direction closer to 90° (Fig S13A), so there was less preview benefit of the fixation in between from previous fixation (Fig S13B) and the duration became longer. However, the CCTELM model fixated longer than subjects when the next saccade nearly reversed the direction (Fig 5C). This was because the previous saccade amplitude was longer for the model under this situation (Fig S13A), and there was less preview benefit (Fig S13B). What’s more, humans and CCTELM model fixated shorter after short and long saccades, but for the CTELM model longer fixations tended to follow large saccades (Fig 5D). Therefore, the CCTELM model explained the subjects’ temporal control of eye movements better.

**Fig 5.**
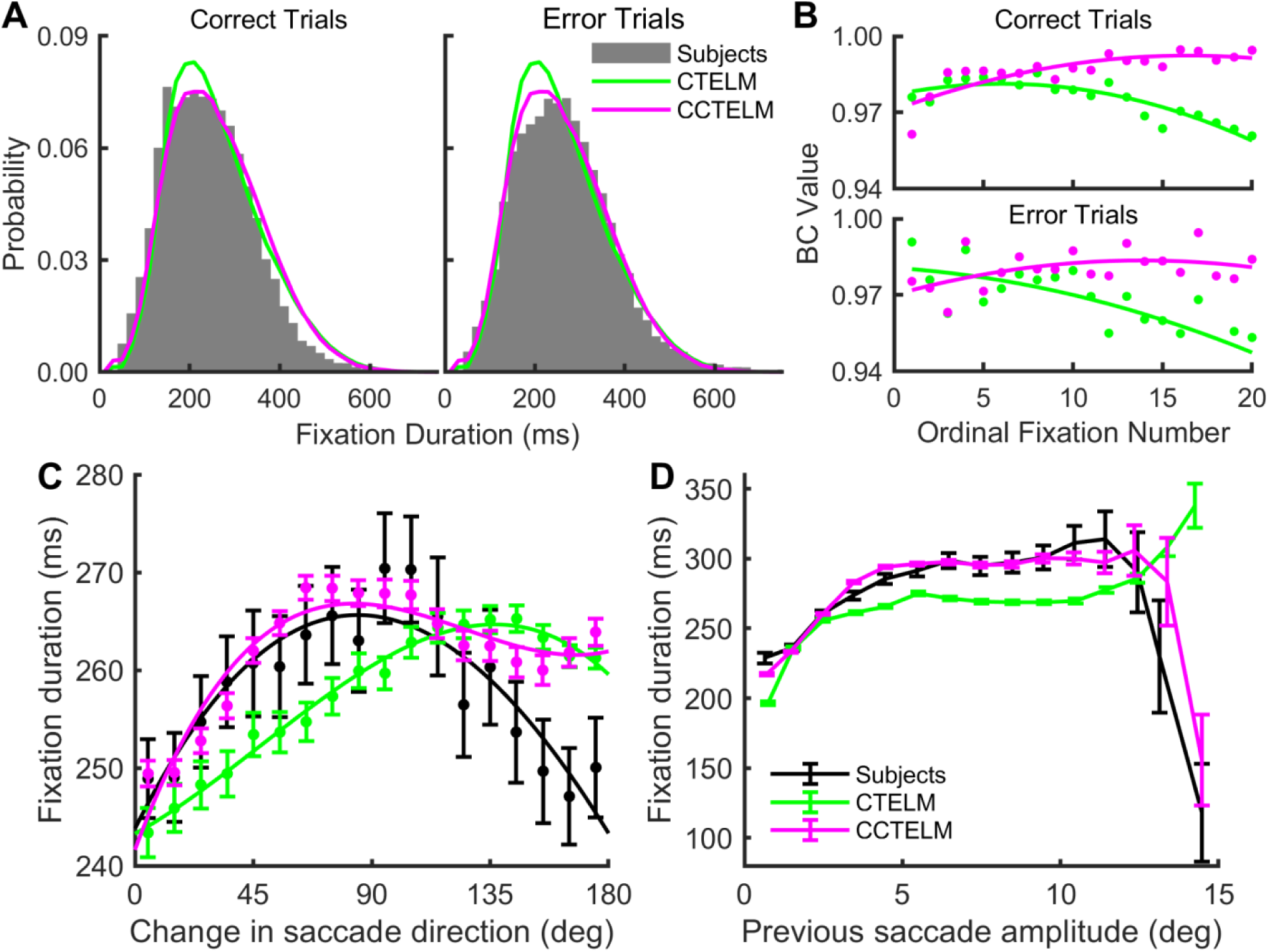
Fixation duration distribution of the subjects and models. **A:** Fixation duration distribution of correct and error trials. The first fixation (when a trial started) and last fixation (when making a response) are excluded. **B:** Bhattacharyya coefficient (BC) between the models’ and subjects’ fixation duration distributions of the initial 20 fixations (first and last fixation are excluded) in correct and error trials. Dots represent raw data; curves represent quadratic functions fitted to the dots. See Fig S11 and Fig S12 for the distribution of raw data at each ordinal position in correct and error trials. **C.** Relationship between fixation duration between two consecutive saccades and the change in saccade direction in all trials. **E.** Relationship between fixation duration previous saccade amplitude in all trials. In C and D error bar represents 95% confidence interval.

The above detailed comparison of eye movement metrics showed that optimal visual search models could only predict a few human eye movement metrics (Fig 3A, Fig 5A). Many other details, especially the overall fixation density and saccade amplitude, the distribution of eye movement statistics as a function of fixation/saccade order, and the dependency between successive eye movements, showed remarkable difference between human and optimal models, and could be well predicted by a model with constraints on saccade accuracy, saccade amplitude and memory capacity. Therefore, the subjects were clearly not using the optimal eye movement strategy in the sense of fully utilizing the visibility map to search the target.

### The Suboptimal Eye Movement Strategy Slightly Impaired Search Performance

Since subjects used a suboptimal search strategy, we next compared the search performance of humans and the three models. With the appropriate choice of target detection threshold *θ_T_* (Table S1), the three models’ correct response rate was similar to the correct response rate of subjects in training set (about 88.0%, see Table 1). The subjects in the testing set had lower correct response rate (84.7%), which may be caused by individual variability and difference in prior training between the two groups of subjects (see discussion).

**Table 1.** Visual search performance of the two groups of subjects (training and testing set) and three models. P(Correct) is the correct response rate.

|  | P(Correct) | Median Fixation Number |  | Maximum Fixation Number |  |
| --- | --- | --- | --- | --- | --- |
|  |  | Correct Trials | Error Trials | Correct Trials | Error Trials |
| Training Set | 87.98% | 8 | 38 | 176 | 158 |
| Testing Set | 84.73% | 10 | 39 | 181 | 425 |
| ELM | 87.31% | 7 | 6 | 42 | 40 |
| CTELM | 88.07% | 7 | 7 | 38 | 36 |
| CCTELM | 88.14% | 8 | 8 | 127 | 96 |

In all correct trials, the ELM and CTELM model searched fastest in terms of median fixation number (7 fixations), followed by the subjects in training set and CCTELM model (8 fixations), and subjects in testing set (10 fixations, see Table 1). The distribution of fixation numbers of subjects and models had a similar peak, but the distribution was more heavy-tailed for human subjects and CCTELM model (Fig 6**Error! Reference source not found.**A, also see maximum fixation numbers in Table 1). With respect to fixation number as a function of eccentricity, the CCTELM model only searched slower than both the ELM and CTELM models when target was distant, and was closer to subjects’ performance (Fig 6**Error! Reference source not found.**B). We also found that the current models could not explain subjects’ search performance in error trials (Table 1), but since most of the trials are correct, so at least the model already agreed well with the majority of experiment data. To summarize, though subjects used a suboptimal search strategy, the performance of this strategy was close to the optimal performance.

**Fig 6.**
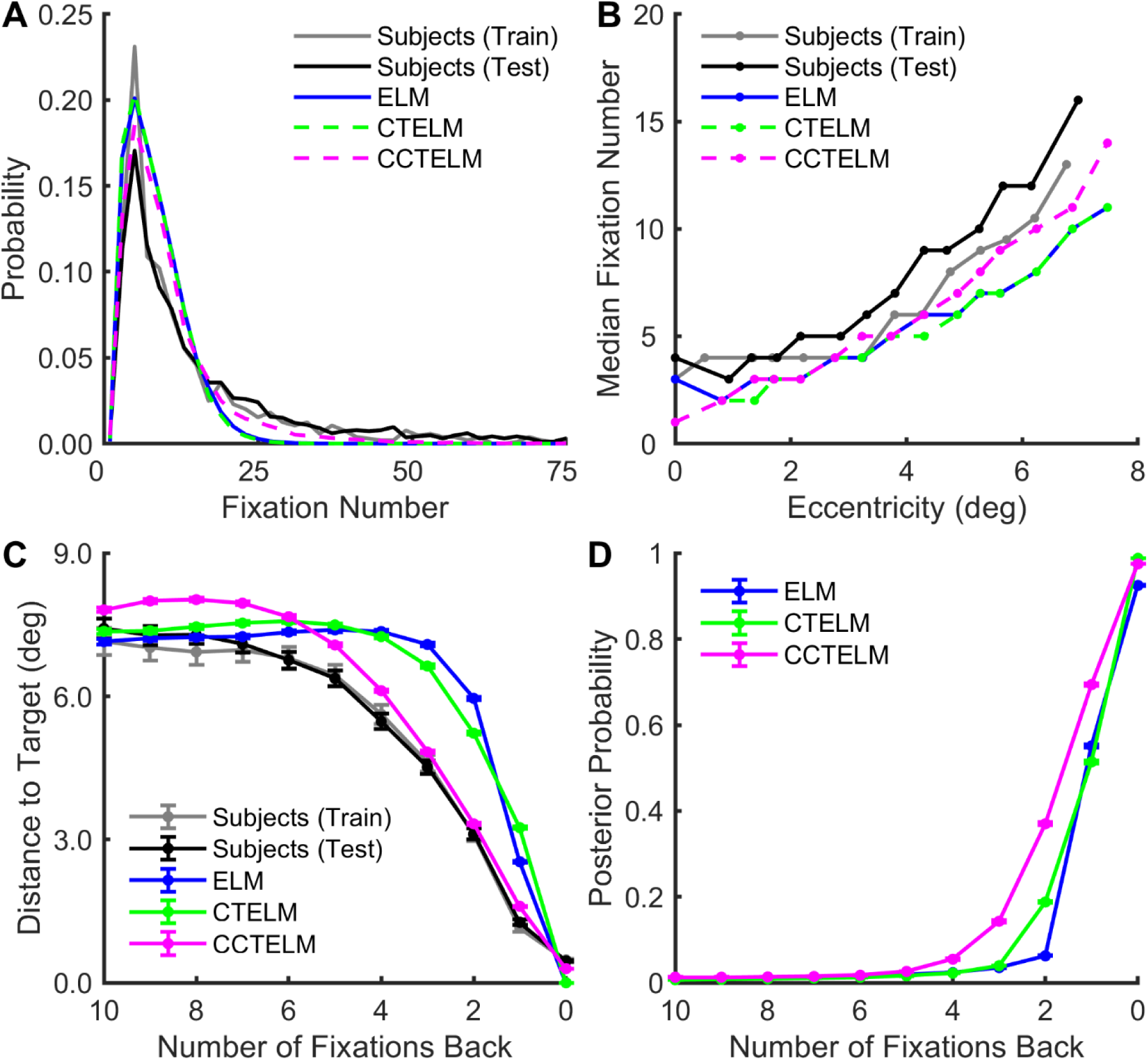
Subjects and the three models’ search performances. **A.** The distribution of the number of fixations needed to find the target in all correct trials. **B.** Median number of fixations to correctly find the target as a function of the median target eccentricity from the image center. **C.** The distance between fixation location and the target location as a function of the number of fixations before correctly finding the target. **D.** Posterior probability at the target location as a function of the number of fixations before the model correctly finding the target. In C and D error bar shows 95% confidence interval.

With the three constraints, the CCTELM model approached the target slower than both the ELM and CTELM models in the last few fixations in a trial, and was closer to subjects’ behavior (Fig 6C). Besides, the posterior probability at target location rose rapidly within the last three fixations for the ELM and CTELM model (Fig 6D), which is similar to the result in [3]. But for the CCTELM model, posterior probability at target location need about five fixations to rise to the decision threshold (Fig 6D). This confirmed that subjects and CCTELM model was less efficient (in the sense of fully utilizing the visibility map) than the ELM and CTELM model though the difference wasn’t large.

As in previous studies [1–3], we measured the visibility map with target location cue in the detection task, one may question the validity of using it in the visual search task when there is no target cue. We also measured the visibility map without a target location cue in four subjects (see methods and S3 Text) and found that the optimal search performance was much slower than the subjects’ performance (Fig S16 in S3 Text). Therefore, in theory, visibility map measured without a target location cue does not allow human-level search performance.

We have shown that the CCTELM model agreed better with human visual search behavior both in eye movement metrics and task performance. The remaining questions were how we estimated the memory capacity, and why these constraints improved the model’s prediction. We have varied the CCTELM model’s memory capacity and examined how the eye movement metrics fit to the training set. Fig 7Error! Reference source not found.A shows that the distributions of fixation distances to screen center and fixation location were most sensitive to memory capacity, and the distributions of fixation duration and saccade amplitude were less sensitive unless at extreme values. On average, the matching to the experimental data peaked when the memory capacity was 8 fixations (Fig 7A). Fig 7**Error! Reference source not found.**B shows the mean number of fixations to correctly find a distant target is affected much more by memory capacity than finding a near target. Together, the results**Error! Reference source not found.** suggest that a model with memory capacity of 8 fixations could significantly improve agreement to experiment data while the search performance is only slightly affected, we therefore choose 8 fixations as the memory of CCTELM model.

**Fig 7.**
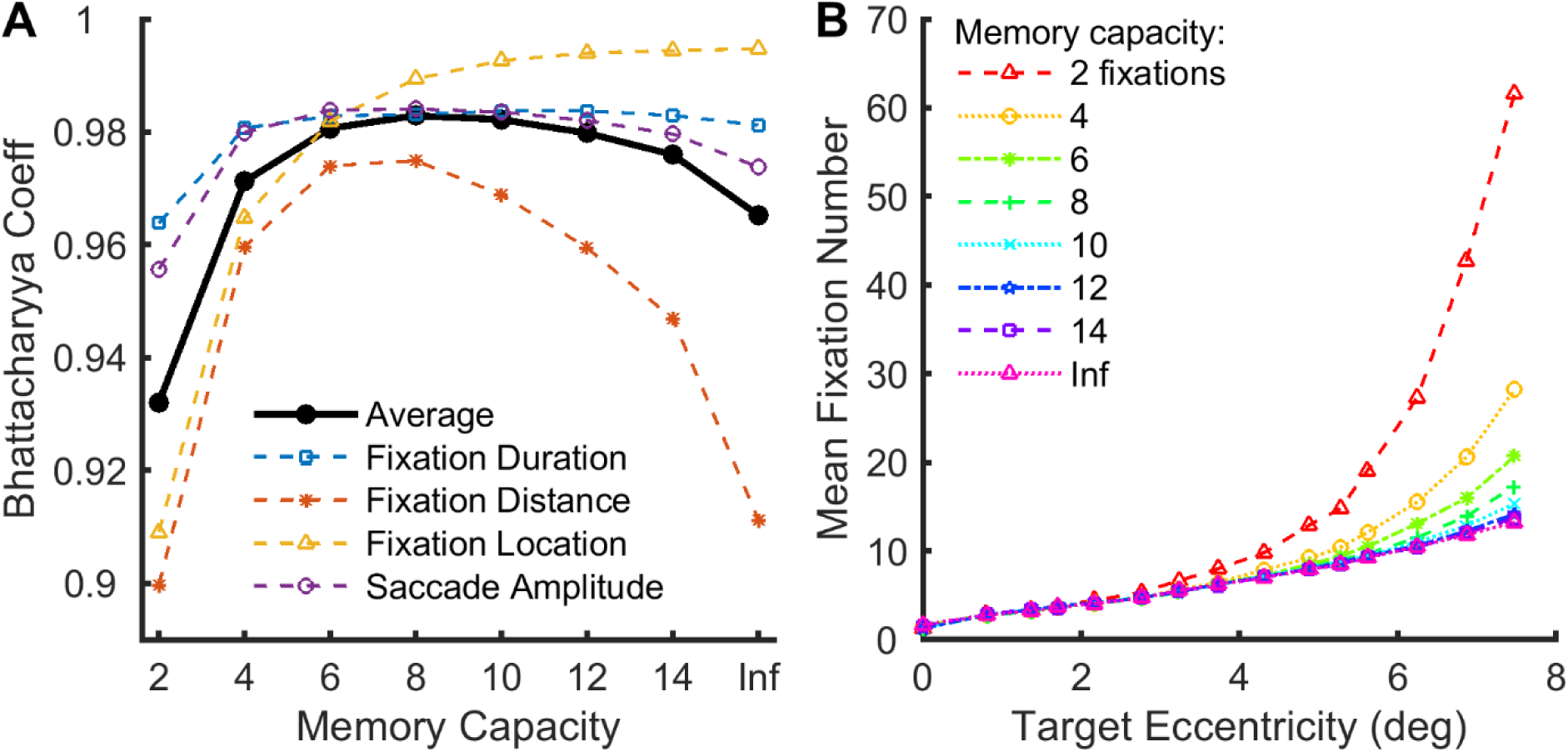
Effect of memory capacity on the CCTELM model’s behavior. **A.** Effect of memory capacity on the goodness of fit to training data. For fixation duration, fixation distance (to screen center) and saccade amplitude, we first align sequences of eye movements at start of the trial and calculated the initial 20 histograms of these metrics from all (both correct and error) trials. Then we calculated the 20 Bhattacharyya coefficients (BC) at each ordinal position and averaged them together to obtain a single value for each memory capacity per metric. For fixation location we calculated the overall 2-dimensional histogram from all trials, smoothed by a Gaussian window with a standard deviation of 0.35 degrees (15 pixels), and calculated the BC value for each memory capacity. The thick black line is the average of the four colored thin dashed lines. “Inf” represents the unlimited memory capacity. **B.** Effect of memory capacity on the mean number of fixations to correctly find the target at different eccentricities of the CCTELM model. The data in A and B came from the CCTELM model whose parameters were fit assuming unlimited memory but evaluated under different memory capacities.

We further checked the scan path of humans and models to understand the effect of saccade amplitude penalty function. Subjectively, the scan path looked similar when the target was close and number of fixations was small; however, difference emerged when the number of fixations was larger (Fig 8). The ELM and CTELM models chose the location that maximize the expected information gain as the next fixation location. In general, one can get most information by fixating at novel regions and avoid previously fixated regions. Therefore, both the ELM and CTELM models tended to avoid image center (where the search starts) and jumped between opposite sides of the search field (Fig 8), producing many long saccades (Fig 4ADE) and larger change in saccade direction (Fig 4C). Adding constraint on saccade amplitude prevent this behavior and forced the model to make series of small saccades around the image to reach the other side of the image (Fig 8), so the change in saccade direction decreased (Fig 4C). This was more similar to humans’ behavior as some of our subjects did report after the experiment that they fixated around the image to find the target.

**Fig 8.**
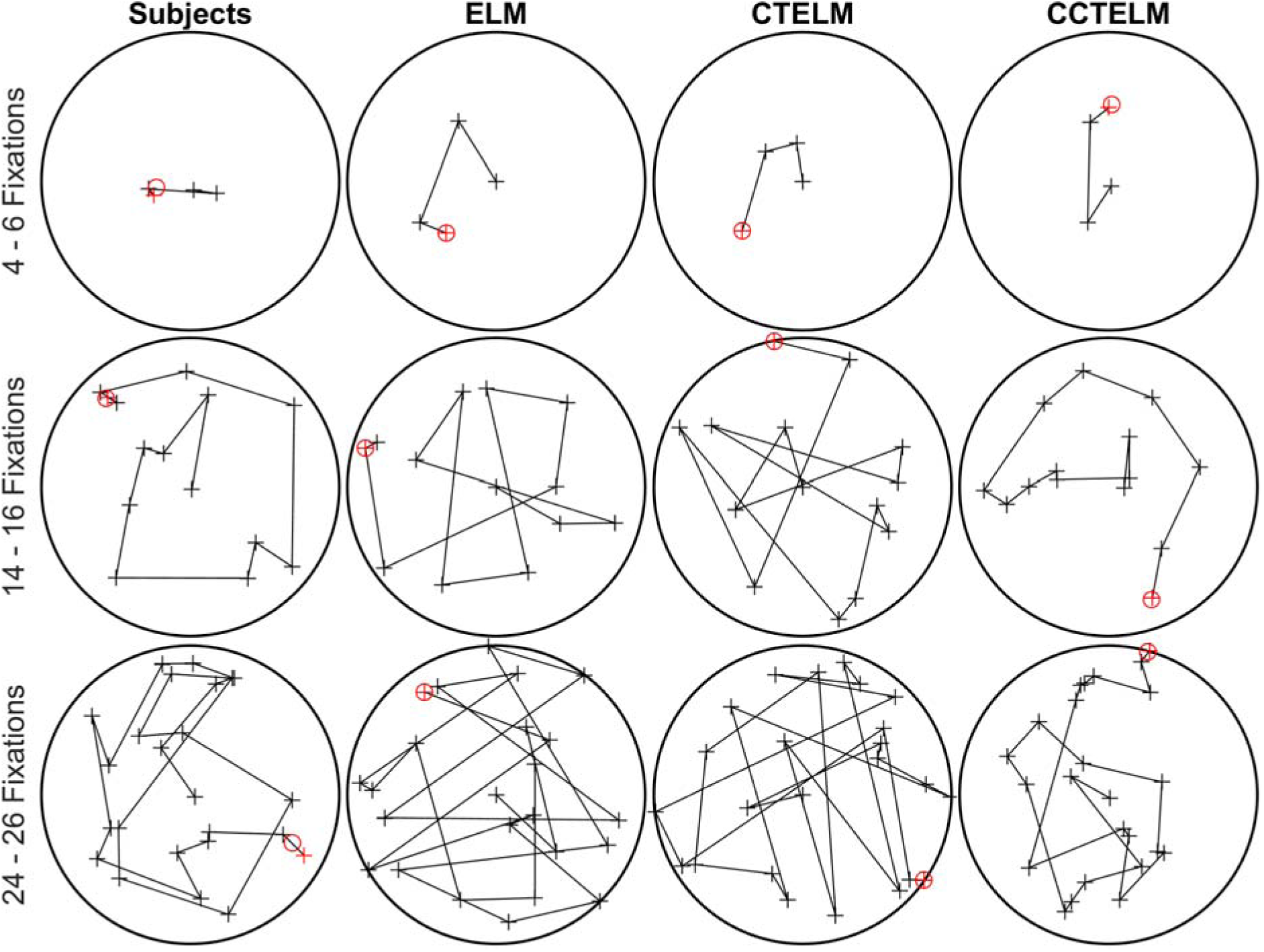
Example scan path of the subjects and models. Scan paths on each row have similar fixation number shown at the left side. The search starts from image center. Black circle represents the image boundary, cross represents a fixation, and the line between two crosses represents a saccade. The final fixation to report the target location is labeled in red. The red circle shows the true target location.

## Discussion

Previous computational models predict that the human visual search behavior follows the optimal Bayesian inference [1–3]. However, both our experiments and simulations confirmed that the human visual search is indeed obeying the suboptimal strategy because of the constraints from memory capacity and saccadic bias. This suboptimal strategy, however, did not greatly affect the overall search performance. Including these constraints into the previously proposed optimal model could significantly improve the model’s prediction of the human eye movement statistics.

### Temporal Course of Visual Processing

An interesting result from the detection task is that central vision processed information faster than peripheral vision. The result agrees with previous findings that scene gist recognition depends more on central vision during the initial 100 ms of viewing [41]. At the neuronal level, the initial central advantage is consistent with earlier finding that the discriminability of neural activities for high contrast gratings peaked rapidly 100 ms after stimulus onset [42], but what makes peripheral vision be slower in our task remains ambiguous. A recent neurophysiological study showed that neurons in the anterior part of inferior temporal cortex responded later to objects presented at peripheral than central visual field [43], but the exact neural mechanisms leading to this phenomenon is still unknown.

One possible reason was that the discriminability of neural activities for suboptimal gratings could peaked much slower [42], and that peripheral vision generally tunes for lower spatial frequencies. From the perspective of our evidence accumulation model, it is possible that the visual system shortens the integration time at central vision to achieve a faster response speed but lengthens the integration time at the peripheral region to compensate for the low signal-to-noise ratio.

### The Control of Fixation Duration

The CCTELM model controlled fixation duration by a drift-diffusion process based on experimentally measured temporal course of visibility and an artificial collapsing decision threshold. Our approach shared some similarities to some previous eye movement models which controlled fixation duration collectively by an autonomous timer and modulation by processing demand from the current fixation [27,28,30]. From this viewpoint, the CCTELM model implemented the autonomous timer as the collapsing threshold, and modulation from the current fixation as the drift-diffusion process. Fixation will be prolonged if the model cannot determine whether the current fixation location has a target because the probability value at the fixation location will not decrease quickly. However, our approach differed from previous models in that the diffusion process could be constructed from experiment data, thus being more biologically realistic, and it is based on probability values so it can be incorporated into the ideal searcher theory. The introduction of a collapsing threshold may seem arbitrary, but its existence, though still controversial, has received certain experimental support [44, 45], and it allows human to trade off accuracy for speed if the current visual information remains ambiguous about target location.

Recent models of eye movement are moving from separate control of fixation location and fixation duration to a more integrated spatiotemporal control strategy [29]. Although the CCTELM model seemed to adopt a separate control mechanism (Fig 2), the decision of fixation duration and fixation location were both based on posterior probability map so they were not completely independent. The model fixated shorter after a smaller saccade (Fig 5D) because part of the information at the new fixation location had already been obtained from the previous fixation, so the probability value at the new fixation location was closer to decision threshold. Besides, long saccades were not accurate, so the model was likely to miss the saccade target and fixated on place with low probability value which made the fixation shorter (Fig 5D). The preview benefit could also explain the relationship between fixation duration and the change in saccade direction (Fig 5C). Though the model fixated longer than subjects when the next saccade nearly reversed the direction, previous study has reported relationship similar to our model’s prediction in visual search task [8].

### Memory Capacity of Visual Search

An interesting discovery from this study is that the subjects’ eye movement statistics predicts the memory capacity in visual search. Similar capacity value has also been used in some models of scene viewing [46], but many models of visual search still ignore this memory limitation. We found that the memory capacity had a considerable effect on model’s eye movement statistics and its agreement to experiment data, so it might be worthwhile for future simulation studies to consider this factor.

One may wonder why the model needed a memory capacity of about 8 fixations when correctly identifying the target needs only 5 fixations before the response (Fig 6D). Memory helps not only in identifying the target, but also in rejecting regions without target so that the model had a higher chance to meet the target in the future, which Fig 6D didn’t show. The simulation experiment with varies memory capacity showed that the model’s prediction on overall fixation location distribution decreased rapidly with less memory capacity (Fig 7A purple dashed line). This was actually because with less memory the model would quickly forget information near the image center (where the search started), and by the ELM rule the model tended to place subsequent fixations near the image center to gain maximal information (fixating near the image boundary causes part of the visibility map to be wasted). Therefore, fixation locations were much more concentrated to the image center in low capacity compared to high capacity model. In other words, larger memory promotes the model to explore outer part of the image, and was therefore needed to efficiently find target at distant locations (Fig 7B).

The type of short-term memory that we modeled is similar to the fragile visual short-term memory (FVSTM). FVSTM is thought to be an intermediate stage between iconic visual memory and visual working memory. It has a high capacity of approximately 5-15 items, lasts for at least 4 seconds [47] and has been shown to be tied to the location of the original visual stimulus [48], making it particularly suitable for integrating information across multiple fixations to generate a posterior probability map. However, the exact memory capacity in visual search tasks remains controversial. There are reports indicating that visual search has a memory of 3 – 4 items [49], 7 – 9 items [50, 51], and more than 10 items [52]. The predicted memory capacity of 8 fixations of the CCTELM model lies within the range of these reported results and thus may serve as additional evidence in this debate. However, the exact memory capacity requires future investigations and may serve as a test of our model.

### Human Visual Search Strategy

The comparison of eye movement statistics between humans and both the ELM and CTELM models showed significant difference on the distribution of fixation location, fixation distance to image center, saccade amplitude, and the dependency between successive eye movements. The CCTELM model further suggested that the discrepancy was mainly due to constraints on saccade amplitude, saccade accuracy and memory capacity. The humans’ preference of short saccades compared to optimal model was also shown in previous study [1]. Therefore, if we consider the degradation of visibility in peripheral vision as the only limitation to visual search, the human eye movement statistics clearly deviated from the optimal model. In this sense, the subjects employed a suboptimal eye movement strategy to find the target.

However, this suboptimal strategy only increased the median fixation numbers by one in the training set and three in the testing set. This relatively small impact, combined with the physiological meanings of the constraints in CCTELM model, suggests that humans were balancing task performance and costs when making eye movements to search for target. If humans try to use the optimal eye movement strategy to fully use the visibility map, they will need to make more long saccades, which means more time spent on saccades [14–16], more subsequent corrective saccades because of decreased landing accuracy [9–13], and stronger saccadic suppression during the saccade [17]; besides, they need to maintain a long history of previous fixation in memory which will also increase the cognitive load. Choosing a suboptimal eye movement strategy that reduces these costs while maintaining a close-to-optimal search performance should therefore be a better solution.

Several previous studies have also shown that human eye movements are suboptimal when optimally performing a task required effortful computation and careful planning [4–6], so subjects may simply follow a good heuristics when searching in these noisy images. The advantage of this strategy is that the computational cost of saccade target selection may be further reduced to selecting a few random numbers, as even the relatively computationally efficient ELM rule requires convolving two maps across the visual field in every fixation. The heuristics may be good enough but not strictly optimal for this particular task. From this perspective, the subjects may be close to optimal balance between search performance and the cost to perform the task.

We found that the subjects’ distribution of fixation location was more uniformly distributed in the search field compared to previously published results on a similar experiment [1, 2]. The subjects in training and testing set also showed some difference on the correct response rate and search performance. This could be caused by both individual variability of visibility map and search behavior and difference of the amount of training prior to the visual search experiment. Previous results in [2] came from two subjects who had measured the target visibility with much more trials at more retinal locations, and had performed more visual search trials with more target visibility levels, so it is possible that previous results were the behavior of two over-trained subjects who performed closer to the optimal model. In our study we had recruited more subjects, so because of time limitation we could not obtain a very densely sampled visibility map and perform many visual search trials, and we did not measure the full temporal course of visibility on some subjects in the testing set before the visual search experiment. This may cause the performance difference between different groups of subjects, but the current results may be more generalizable to a larger pool of subjects.

### Limitations of Current Model

Although the CCTELM model explained a range of behavioral data of visual search, it is still a simple model and did not fully reflect the complexities of the visual system. First, accurately measuring the target visibility is challenging, because the detection task may not reproduce the actual attention deployment and pre-saccadic enhancement of visual processing [53] happened in visual search, and introduced perceptual learning at measured locations. Second, humans may plan multiple fixations ahead during a fixation [54–56], whereas our model only plans one fixation ahead. Third, the implementation of memory is still relatively simple. The model either kept information from a fixation or completely forgot it, whereas human brain memory may gradually decay over time [57, 58], and this was considered in previous model of scene viewing [46]. In addition, multiple types of memory may be involved in visual search [59, 60], but this was not reflected in the current modeling approach.

Subjects also had some eye movement features that the CCTELM model could not reproduce. First, human saccade direction is biased toward horizontal and to a less extend toward vertical directions [46, 61]. The previous optimal models of visual search showed that the horizontal bias could be explained by an horizontally elongated visibility map measured in experiment [1, 2]. However, our visibility map data is more radially symmetric (Fig 1E) so we did not see the same modeling result as previous studies. There are probably other factors (such as the distribution of features in natural scene) leading to this bias in humans [61], and are not captured in the constraints of the CCTELM model. Second, the model did not spend as many fixations as subjects did in error trials. One possible reason is that we used the dynamic noise instead of the static noise which considers the variability of task difficulty due to the noisy image background superimposed on the target. But since the target is well visible under the chosen contrast (*d’* = 3), the model with dynamic noise already agreed with correct trials, and previous study found only small effect of static noise on the optimal model’s behavior [2], we chose to use dynamic noise in this study for its computational simplicity. We hope that this model could help developing more complete models of eye movement in the future.

## Materials and Methods

### Human Subjects

Ten student volunteers (6 males and 4 females, age range 19-30 years) participated in the experiment for monetary compensation. One subject was the author (Y.Z). All subjects had normal or corrected-to-normal vision.

### Ethics Statement

This study was approved and supervised by the Ethics Committee of the School of Life Sciences at Fudan University (BE1942), and written informed consent was obtained from all subjects.

### Apparatus

Stimuli were presented on a 24.5-inch BenQ ZOWIE XL2540 LCD monitor at a resolution of 1920 × 1080 and a refresh rate of 240 Hz. Subjects were seated 70 cm in front of the display with their heads fixed by a forehead and chin rest. Stimuli were generated and presented using MATLAB (MathWorks) and the Psychophysics Toolbox extensions [62–64] running on a Windows 7 (Microsoft) machine. Eye movements of both eyes were recorded at 2000 Hz with a TRACKPixx3 eye-tracker (Vpixx Technologies, Canada).

### Experiment Design and Statistical Analysis

#### Target and background image

The target was a six cycle-per-degree Gabor with a diameter of 0.3 degrees, oriented 45 degrees counterclockwise from the vertical, and windowed by a symmetrical raised cosine (half-height width was one cycle of the Gabor). The background image was a circular naturalistic noise image (whose statistics follow a 1/*f* power spectrum) placed at the center of the screen with a diameter of 15 degrees and root-mean-squared (RMS) contrast of 0.2 (Fig 1A). The RMS contrast was defined as the standard deviation of the pixel gray value divided by the mean gray value. The area outside the background image was set to the mean gray value of the image (0.5). The background images used in each trial of detection task and visual search task were randomly picked from a database of 1000 independently generated images.

#### Eye-tracker calibration

During each laboratory visit, subjects first performed a 13-point calibration routine. The outer edge of the 13 points formed a square that covered the circular search area. We repeated the calibration until the average test-retest measurement error of both eyes fell below 0.5°. Recalibration of the eye-tracker was performed whenever the eye-tracking accuracy failed to meet the requirements of the experiment (described below).

#### Selecting target RMS contrast

In this experiment we measured the relationship between foveal target visibility and target RMS contrast and selected the target contrast level that made the foveal visibility equal to 3.0. Each participant therefore had a unique target contrast and was used for all subsequent experiments. The goal was to normalize the target detection difficulty across subjects.

At the start of each trial, a black cross and stroked circle appeared at the center of the screen, indicating the fixation location and target location. The remaining display area was set to a 0.5 gray value. Subjects began a trial by fixating within 1° from the screen center and pressing a button. The cross and circle then disappeared for 200 − 400 ms (chosen randomly), followed by two 250 ms intervals of the stimulus image separated by a 500 ms gray interval. The two stimulus intervals had the same noise background image; however, a randomly chosen stimulus had a target at the screen center. The subjects then chose the interval that contained the target with a keyboard. A trial was aborted if the participant fixated more than 1^°^ away from the screen center at any time during the trial. This section contained 4−5 levels of target contrast (depending on the participant’s pace), and each level included 200 trials. Trials with different target contrasts were randomly interleaved. We chose the appropriate target contrast levels (ranging from 0.03 to 0.12) by a pilot experiment for each participant to avoid floor or ceiling effects.

We first calculated the hit rate and correct rejection rate of each target contrast level from behavior data. The hit (or correct rejection) rate was defined as the number of trials that target appeared in the first (or second) interval, and the participant made a correct response, divided by the total number of trials that target appeared in the first (or second) interval. The relationships between the hit/correct rejection rate and target RMS contrast *C* were then separately fit with a Weibull function:

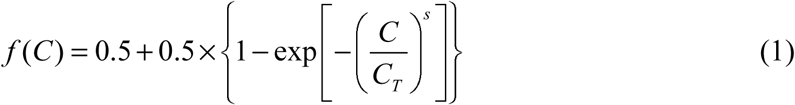

The threshold parameter *C_T_* and steepness parameter *s* were estimated by maximum-likelihood methods implemented in the Palamedes Toolbox [65]. The relationship between the foveal target visibility and target RMS contrast could then be estimated by signal detection theory [66]:

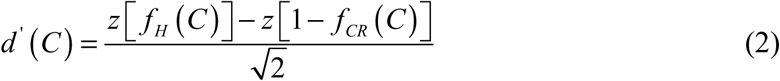

Here, *z*(*x*) is the inverse standard normal cumulative distribution function. *f_H_*(*C*) and *f_CR_*(*C*) are the estimated hit and correct rejection rates given by Equation (1). We could then obtain the target contrast level that made *d^’^*(*C*) *=* 3.0.

#### The detection task

The goal of the detection task was to measure the change of target visibility over time in the visual field. The experiment had three versions. Four subjects participated in the first version, four participated in the second version (the author Y.Z participated in both), and three participated in the third version.

In the first and second versions we measured the relationship between the stimulus presentation time and target visibility along four cardinal directions in the search field. The procedure was the same as the experiment to select target RMS contrast, except that we varied both the stimulus duration and target location. In the first version of experiment, subjects were required to fixate at the screen center and decide which of the two stimulus intervals contained the target at the cued location. The experiment contained five blocks (four cardinal directions plus the screen center). For each block, we chose 3−4 different target locations (except when the target was at the screen center) and 4−5 levels of stimulus presentation time. The target was located 0−6.75° (0−90% of the image radius) away from the screen center. When the target appeared at the screen center, the stimulus duration was chosen from 4 to 200 ms, and each level was tested for 100 trials. When the target appeared at peripheral locations, the stimulus duration was chosen from 50 to 700 ms, and each combination of stimulus length and target location was tested for 50 trials. In the second version of experiment, the target location cue was not shown, and in each trial, the target location and stimulus presentation time were randomly chosen from all possible combinations in the first version of experiment. The results of the second version are shown in Fig S15 in S3 Text, but it was not used in the results of the main text. The third version was the same as the first version except that there is only one level of stimulus presentation time (250 ms). It was used to measure the visibility map efficiently and get enough training to prepare for the visual search experiment, so the data from this version was not used in this manuscript.

The first and second version of the detection task contained about 5000 trials. Subjects finished these trials by multiple laboratory visits within two months. Before the start of the formal experiment, they practiced with 25-100 trials to stabilize the performance. Subjects received feedback about their performance during practice trials but did not receive feedback during the formal experiment.

#### The evidence accumulation model

Behavior data from different subjects in the first and second version of the detection task was first separately pooled together. Target visibility at each location for each level of stimulus duration was then calculated by Equation (2). The data was used to formulate an evidence accumulation model that described the change of target visibility over time, and this model was later used in the continuous-time eye movement model. According to signal detection theory, visibility is essentially the distance between noise and signal-plus-noise distributions. Therefore, we extended the template response model to visual stimulus at each location in the visual field used in [1–3] to a drift-diffusion process *W_t_*.

Suppose the visual evidence Δ*W* generated at one location within a short time interval Δ*t* is independently drawn from a normal distribution. If the target appears at this location, the distribution is *N*(Δ*t,*σ^2^Δ*t*), otherwise *N*(−Δ*t,*σ^2^Δ*t*). The value of standard deviation σ controls target visibility at this location. If the evidence at each time step Δ*t* is integrated perfectly, then the total evidence at time *T* = *n*·Δ*t* follows the distribution of the sum of *n* normal random variables:

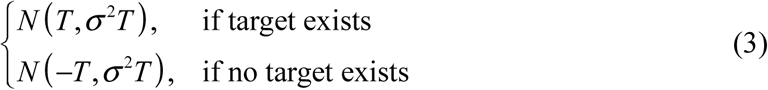

According to signal detection theory [66], the evolution of target visibility *d^’^* over time is:

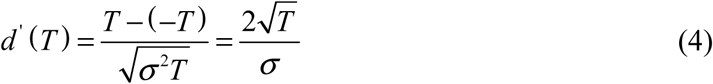

Equation (4) implies that target visibility will increase to infinity at any retina location given unlimited amount of viewing time, which is impossible. This problem can be resolved if we assume that the weight *w* of previous evidence decays exponentially by a speed parameter *k* over time:

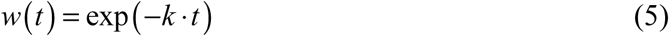

Suppose the total accumulated evidence at one location from 0 to the *i^th^* time step is *W_i_* (*W_0_ =* 0), then the by the (*i* + 1)*^th^* time step, the total accumulated evidence is:

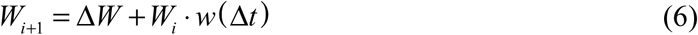

Then the total acquired evidence *W_n_* at time *T* = n·Δt follows the distributions:

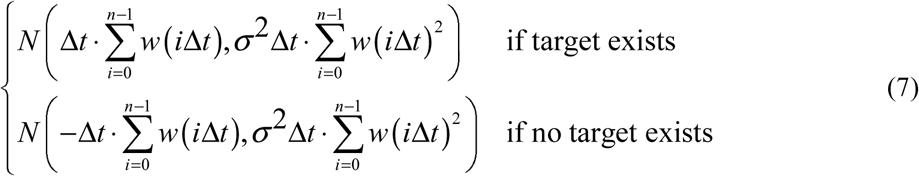

Note that:

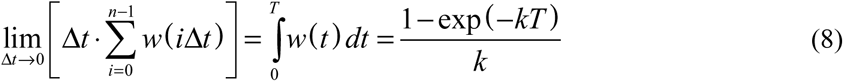

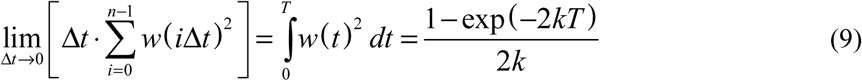

Substituting Equations (8) and (9) into Equation (7), the evolution of target visibility over time can be formulated as:

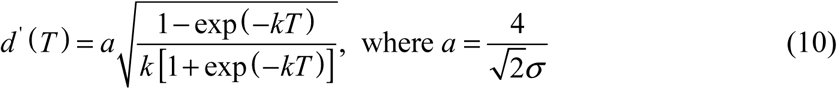

We first fit Equation (10) to the temporal course of target visibility at each measured visual field location (Fig 1E) and got a series values of parameters *a* and *k* at different locations (*x*, *y*) relative to the fixation location (0, 0). We found that the values of these two parameters decay exponentially from fixation location (Fig S5), so a simple function describing parameter *a* (or *k*) is:

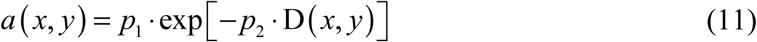

Here 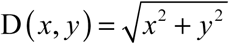 is the distance between measured location to fixation location. To capture potential asymmetry between horizontal and vertical visibility, Equation (11) is further modified to be:

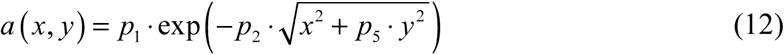

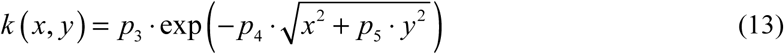

Equations (10), (12) and (13) together describe the change of visibility map over time. The parameters *p*_1_, *p*_2_, *p*_3_, *p*_4_, *p*_5_ in these equations were estimated by minimizing the MSE between the calculated and measured visibility data using the GlobalSearch algorithm in the Global Optimization Toolbox of MATLAB.

#### The visual search task

In this experiment, subjects were asked to search for the target within the background noise image as fast as possible while trying to maximize the response accuracy. The target locations were sampled uniformly inside the search area, and subjects were informed about this feature. Subjects began each trial by fixating within 1° away from a cross displayed at the screen center and pressing a button. Then, the cross disappeared for a random interval (0.2-0.3 seconds), and the search image appeared and waited for subjects’ responses. There was no limit in waiting time. When the subjects found the target, they fixated at the target and pressed a key on the keyboard. The final fixation location and the target’s true location were then shown on the screen. If the distance between these two locations was less than 1°, we labeled the response as correct. Subjects were asked to report if they had found the correct target location, but the response was judged as incorrect. In this case we would recalibrate the eye tracker. Subjects performed 200−500 (mean = 315) visual search trials depending on their available time.

#### Eye-tracking data analysis

Eye-tracking data were analyzed by the EYE-EEG extension [67] of the EEGLAB toolbox [68]. We used the algorithm described by Nyström & Holmqvist (2010) to classify eye movements into saccades and fixations. Inaccurate eye movement events from noisy period of one eye were replaced by events from the other eye. Fixations separated by blinks were recognized as two fixations. Data from the four subjects in first version of detection task were merged and served as training set to fit the parameters in eye movement model (described below), and data from the rest six subjects were merged and served as testing set. The training set contained 18941 fixations and 19603 saccades from 1190 visual search trials, and the testing set contained 39985 fixations and 40010 saccades from 1958 visual search trials.

When summarizing fixation durations, we excluded the first (when subjects fixated at screen center) and the last fixations (when a response was made) to avoid the effects of the anticipation and motion preparation on the fixation duration. When summarizing fixation locations, we excluded the first fixation at the screen center.

### The Entropy Limit Minimization (ELM) Model

We used the ELM model [1] as our baseline model because it greatly reduced the computational burden compared to the Bayesian ideal model while maintaining a similar search performance [1, 3] and has received electrophysiological support [70]. The model could only fixate at 400 locations uniformly sampled inside the search field (Fig S6). We set the fixation duration to 250 ms and evaluated the visibility map using Equation (10). The visual information *W_i_*_,*LF*_ obtained at location *i* (*i* = 1 *n*, *n* = 400) during the *F^th^* fixation at location *L_F_* was sampled from normal distribution 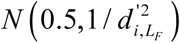 if the target was at this location, otherwise from normal distribution 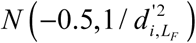. The ELM model did not model the distribution of fixation duration, so the visual information at each location was sampled once per fixation. The posterior probability map of target location was calculated from information gathered in all previous fixations (indicating unlimited memory) according to Bayesian theory [1]:

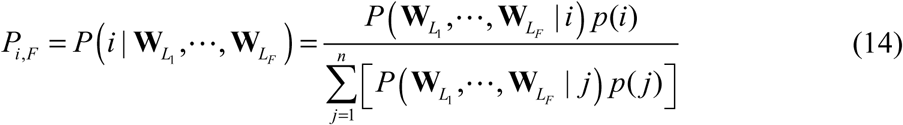

Here *p*(*i*) = 1/*n* is the prior probability of the target being at location *i*. The vector 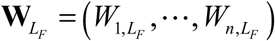 is the visual evidence at all locations gathered during the *F^th^*fixation at location *L_F_*. Assuming the noise was independent at each location and during each fixation, we have:

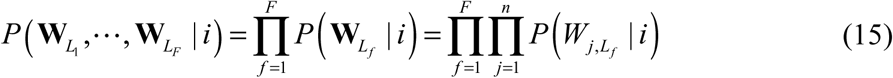

Since *W_j_*_,*L*_ conditioned on target location *i* follows normal distribution 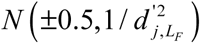 by substituting its probability density function into Equations (14) and (15) we could derive:

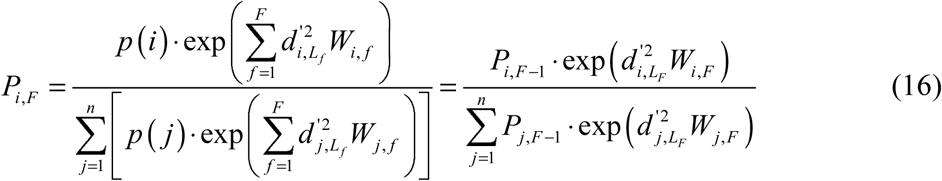

After calculating the posterior probability map of target location, the ELM model selected the next fixation location to maximize the expected information gain (which is the expected reduction of entropy) of the next fixation. Najemnik & Geisler [1] showed that it could be calculated in a very simple form:

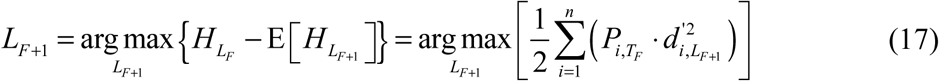

where 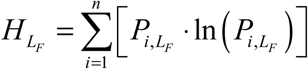 is the entropy of posterior probability of target location.

The model terminated the search whenever the posterior probability of the target being at the current fixation location exceeded a threshold *θ_T_*, and the target location was set as the final fixation location. The value of *θ_T_* was chosen to make the model’s correct response rate comparable to the subjects’ data (see Table S1).

### The Continuous-time ELM (CTELM) Model

The CTELM model proposed here extends the ELM model to account for the distribution of fixation durations. A schematic diagram of the CTELM model is shown in Fig 2 and will be described in the following sections.

#### Temporal course of evidence accumulation

The accumulation of visual evidence over time can be described by the evidence accumulation model discussed in previous sections. In practice, for each fixation *f*, we first calculated the values of parameter 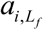 and 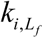 at the 400 locations across the visual field according to Equations (12) and (13). These values were calculated only once per fixation. Then for each time step Δ*t* (1 ms), a random evidence sample 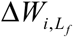 was generated at each location *i* from the normal distribution 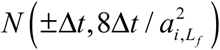 (see the relationship between *a* and σ in Equation (10)) depending on whether target was present at this location. The current total accumulated evidence 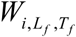 at location *i* at time *T_f_* since the start of the *f^th^* fixation was calculated from 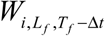 in the previous time step by:

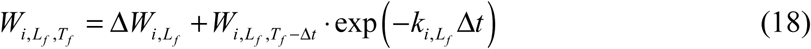

#### Calculating target’s posterior probability map

At each time step, the posterior probability of target being at the *i^th^* (*i* = 1 …*n*, *n* = 400) location could be calculated from the accumulated evidence from the current and all previous fixations by Bayesian theory:

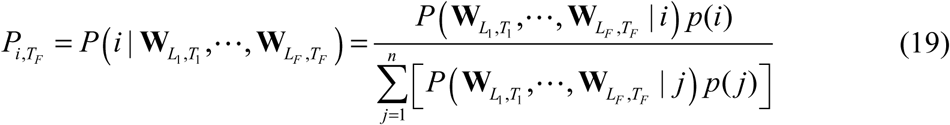

The vector 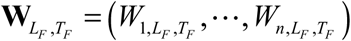 is the accumulated evidence by time *T_F_* of the *F^th^* fixation at all locations. Like the ELM model, the CTELM model also had unlimited and perfect memory across fixations.

Assuming that visual information was independently accumulated at each location during each fixation, Equation (19) could be simplified to (see the S1 Text for details):

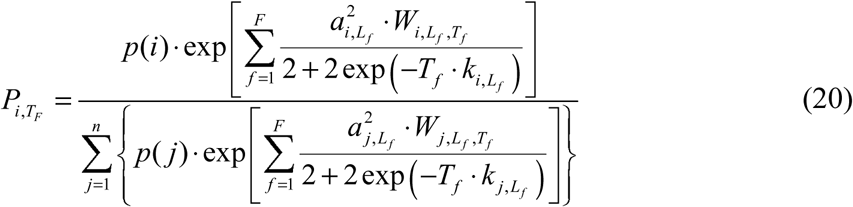

By applying Equation (20) to all target locations at each time step, we could calculate the change in the posterior probability map of the target location over time.

#### Decision on saccade timing

In the CTELM model, saccades were triggered randomly in two different ways according to their relative frequency. The majority (97%) of saccades (normal saccades) were triggered by a decision process concerning whether the current attention location contained the target. The first saccade in a trial was always a normal saccade. If the probability of the target being at the current attention location was lower than a threshold, the model makes a saccade decision. We defined the current attention location as the current fixation location when the next saccade decision had not been made, and as the next fixation location when the next saccade decision had been made, but the eyes had not moved because of the delay from the cortex to the eye movement muscles (saccade lag). We implemented the saccade threshold *θ_S_*(*t*) as a “collapsing bound”, similar to certain decision-making models [44,71–73]:

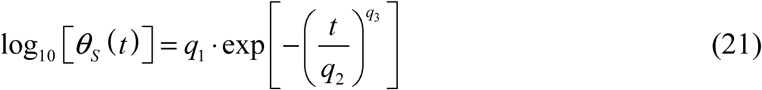

Here, *q*_1_, *q*_2_, *q*_3_ (*q*_1_, *q*_3_ < 0, *q*_2_ > 0) are parameters whose values are estimated by fitting to distributions of fixation durations from experimental data. After the saccade decision was made, a new saccade decision process started immediately.

In addition to normal saccades, a small group of low-latency saccades can be generated by other visual pathways [74, 75]. In the model, we set the frequency of low-latency saccades to 3% according to the frequency of express saccades in an overlap task [76]. The latency (in seconds) is directly drawn from a gamma distribution with shape parameter 10 and scale parameter 6.9 × 10^-4^, plus a delay of 0.03 seconds. This latency follows a distribution with mean = 99 ms, 5% percentile = 68 ms, and 95% percentile = 138 ms, which roughly falls in the 80-120 ms range of express saccade latency [77]. After the saccade decision was made, a new saccade decision process started immediately before the delay of 0.03 seconds.

In the model, the delay from the retina to the visual cortex (eye-brain lag) was set to 0.06 seconds [78], and the saccade lag was set to 0.03 seconds [79]. At the start of a new fixation, the model continued to accumulate information from the previous fixation until the arrival of new information after the eye-brain lag. We did not consider the duration of a saccade itself (a saccade is finished instantly) because of saccadic masking [17].

#### Decision on saccade target

When making a saccade decision, the CTELM model used the ELM rule (Equation (17)) to select the next fixation location. However, the deviation of Equation (17) as shown by Najemnik and Geisler [1] requires the visibility 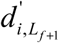 to be a fixed value within a fixation, but in the CTELM model, 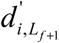 changes with time. As a simplification, we set the expected duration of the next fixation to be 0.25 seconds when evaluating Equation (17). We ran Monte Carlo simulations and found that the expected information gain calculated in this way had a high Pearson correlation (mean = 0.923) with the actual information gain (Fig S14 in S2 Text).

#### Decision on target location

The search stopped whenever the posterior probability at the current fixation location exceeded a target detection threshold *θ_T_*, and the model set the current fixation location as the target location. The value of *θ_T_* was chosen to make the model’s correct response rate comparable to the subjects’ data.

### The Constrained-CTELM (CCTELM) model

The schematic diagram of the CCTELM model is the same as the CTELM model (Fig 2), but we limit the memory capacity, suppress the probability of making long saccades, and add saccade landing position variability to the model.

#### Memory capacity

For simplicity, we set the memory of the model to be all-or-none. The model either retained complete information from a fixation or completely forgot it. A model with a memory capacity of *M* could only integrate information from the past *M*−1 fixations plus the current fixation, so at the *F^th^* fixation, the earliest fixation that the model could integrate was *f_s_* = max(1, *F* − *M*+1). The posterior probability target being at location *i* could be calculated as (see S1 Text for the derivation of this formula):

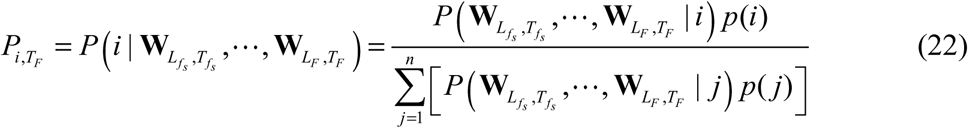

In S1 Text we showed that the expression could be simplified to:

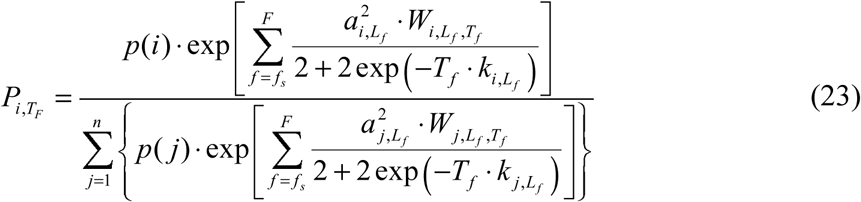

#### Decision on saccade timing

The CCTELM model used the same method to decide saccade timing as the CTELM model. However, because of the saccadic inaccuracy, the CCTELM model could fixate at every possible location in the search field (not just the predefined 400 locations). The posterior probability of target being at current attention location, therefore, was the sum of posterior probability at all the predefined 400 locations within 0.5 degrees away from the actual attention location. Our Monte Carlo simulation showed that in 44.46%, 47.13% and 8.41% cases, one, two, and three nearby locations were counted, respectively.

#### Decision on saccade target

As we show in result section, the ELM and CTELM model made too many long saccades compared to humans. The preference of small saccades on humans has been observed in previous studies [23, 61], and several possible reasons for this preference have been proposed [61]: (1). short saccades decrease the time spent on saccades [14–16]; (2). short saccades decrease the degree of saccadic suppression of motion perception [17]; and (3). short saccades are more accurate and decrease the chance to make a secondary saccade to the target [9–13]. We considered these factors as possible costs to make a saccade, represented by a penalty function H(*L_f_*, *L_f_*_+1_) when selecting the next fixation location:

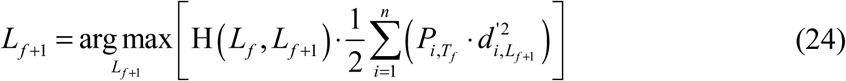

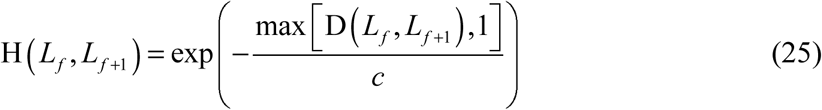

where D(*L_f_*, *L_f_*_+1_) is the Euclidean distance between the current and next potential fixation location in visual degree, and *c* is a free parameter. Thus, even though a distant location may have high expected information gain according to the ELM rule, Equation (25) suppresses the probability making one long saccade to that location and promotes making several short amplitude saccades. We chose not to apply the penalty function within 1 degree away from fixation location to prevent interfering with the inhibition of return phenomenon generated by the model.

#### Variability of saccade landing position

The accuracy of saccades can be described by the offset between mean saccade landing positions and the saccade target, and the variability of saccade landing positions. Some studies found that the mean saccade landing positions were quite accurate for the saccades typically made in our experiment [12,15,16], but some others showed undershooting bias of long saccades (> 7°) [10], or both short and long saccades [13]. However, many existing studies showed that the standard deviation (SD) of saccade landing positions increased linearly with saccade amplitude to a similar scale [10,12,13]. Therefore, for simplicity we only considered saccade landing position variability in the model and assumed the mean landing positions were accurate. We further assumed the actual saccade landing positions formed a 2-dimensional Gaussian distribution around the mean [12], and ignored any difference in horizontal and vertical directions.

To accurately describe the relationship between SD of saccade landing positions and saccade target eccentricity, we analyzed the raw data from experiment 2 of [12]. In brief, this experiment required subjects to fixate at a target with eccentricity varying from 0.5 to 17.5 degrees left or right away from the initial fixation location, and identified the number of small dots presented around the target location [12]. Only the first, non-anticipatory saccades after presentation of the target were analyzed. Raw data was shown in Fig S7A. Leftward and rightward trials with the same eccentricity were combined (leftward saccades flipped to rightward). The SD of saccade landing positions σ_s_ was calculated as:

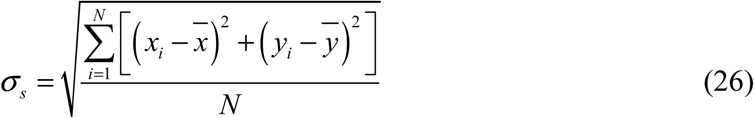

Here (*x_i_, y_i_*) is the saccade landing position of the *i^th^* trial, 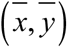 is the mean saccade landing position of all *N* trials with the same target eccentricity. The relationship between target eccentricity and σ_s_ was fit by linear regression (Fig S7B). The values of slope (*m*) and intercept (*c*) coefficients are shown in Table S1.

#### Decision on target location

The CCTELM model stopped searching whenever the posterior probability at the current fixation location (sum of all 400 predefined locations within 0.5 degrees from fixation location) exceeded a target detection threshold *θ_T_*. The model set the target location as the predefined location with highest posterior probability.

The value of *θ_T_* was chosen to make the model’s correct response rate comparable to the subjects’ data.

### Fitting Model Parameters

Table S1 shows a summary of the parameters used in the ELM, CTELM, and CCTLEM model. All the parameters were derived by fitting the models to the data of training set (4 subjects, including both correct and error trials).

For the CTELM model, there are three free parameters in Equation (21) that control the distribution of the fixation duration. The remaining parameters were estimated from experimental data and fixed when fitting the free parameters. The values of the free parameters were estimated by genetic algorithm [80] that minimized the sum of the Bhattacharyya distance (DB) between the simulated and experimental histograms of the fixation duration (*T_sim_*, *T_exp_*):

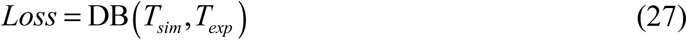

where the *DB*(*H*_1_, *H*_2_) of the two histograms *H*_1_ and *H*_2_ is defined as

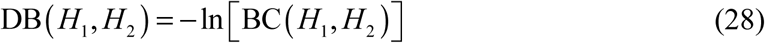

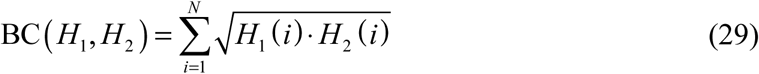

Here, *N* represents the number of bins in the two histograms. The histogram should be calculated as probability. The genetic algorithm was implemented in the Global Optimization toolbox of MATLAB. We set the population size to 150 and max generations to 25, leaving the other parameters unchanged. For each parameter sample in the population, the model was run for 1000 times to obtain the histogram of fixation durations. We used the Multicore package (https://www.mathworks.com/matlabcentral/fileexchange/13775-multicore-parallel-processing-on-multiple-cores) to run the genetic algorithm on multiple computers.

For the CCTELM model, we fit the four free parameters in Equations (21) and (25) by genetic algorithm. The cost function is the sum of the Bhattacharyya distance (DB) between the simulated and experimental histograms of the fixation duration (*T_sim_*, *T_exp_*) and saccade amplitude (*A_sim_*, *A_exp_*):

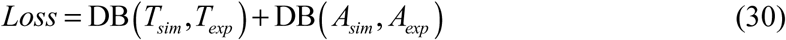

We set the population size to 250 and max generations to 25, leaving the other parameters unchanged. For each parameter sample in the population, the model was run 1000 times to obtain the histograms of the fixation durations and saccade amplitudes.

Recently there are proposals to find free parameters in eye movement models by maximum likelihood estimation (MLE) [81]. Indeed, MLE has many merits over the genetic algorithm that we used here, but it requires getting the likelihood function which is hard for complex models like the CCTELM model. An alternative and popular likelihood-free method, called the Approximate Bayesian Computation (ABC), works by evaluating the model with different combinations of parameter values and minimizes the distance (by calculating certain summary statistics) between simulated and experiment data [82].

However, with only a few parameters the speed of ABC gets very slow because of the large parameter space to explore [83]. Therefore, we chose to use the genetic algorithm to estimate free parameters which has been used in previous eye movement models [27,28,84]. Although it is not mathematically well-founded compared to MLE or ABC, in practice we found that the method consistently produced good fitting to experiment data in reasonable amount of time.

To find the memory capacity that best predicts the experiment data, we first assumed unlimited memory capacity and fit the four free parameters to subjects’ data. We then evaluated the model with memory capacity of 2, 4, ⋯, 14 and infinite fixations for 50000 times each. The value of target detection threshold *θ_T_* was set to 0.952 (Table S1). For the simulation result belonging to each capacity, we calculated the histograms of fixation duration, fixation distance and saccade amplitude of the initial 20 fixations and saccades, and the overall distribution of fixation location. Both correct and error trials were used. Then for fixation duration / distance and saccade amplitude, we calculated the Bhattacharyya coefficient (BC) between histograms of simulation and experiment data corresponding to the same fixation or saccade ordinal index. Thus, for each memory capacity value, we obtained 1 BC value for fixation location and 20 BC values for fixation duration, fixation distance, and saccade amplitude. Finally, the 20 BC values were averaged to obtain a single BC value per eye movement metric.

### Simulating Visual Search

The three models were evaluated for 50000 trials to obtain eye movement statistics. We randomly rotate the 400 potential target locations around the image center before each trial to help obtain a smooth distribution of fixation locations. The target locations were randomly selected with equal probability in each trial.

When we summarized the eye movement metrics, we excluded the first and the last fixations for the fixation duration, and excluded the initial fixations for the fixation location. We used BC to quantify the similarity between the histograms of the model’s and humans’ eye movement metrics. When the histograms are calculated as probabilities and have the same number of bins, BC has a maximum value of 1 if the two histograms are completely the same and a minimum value of 0 if the two histograms have no overlap.

### Data and Code Accessibility

The experiment and simulation results, the code for analyzing eye-tracking data and visual search models are available on Open Science Framework at https://osf.io/ypcwx/. Questions should be directed to the corresponding author.

## Acknowledgments

We thank Professor Zhaoping Li for her advice on the manuscript.

## Supporting Information

Fig S1. Temporal course of target visibility at four cardinal directions (shown in four subplots) relative to fixation location of subject 1 in training set.

Fig S2. Temporal course of target visibility at four cardinal directions (shown in four subplots) relative to fixation location of subject 2 in training set.

Fig S3. Temporal course of target visibility at four cardinal directions (shown in four subplots) relative to fixation location of subject 3 in training set.

Fig S4. Temporal course of target visibility at four cardinal directions (shown in four subplots) relative to fixation location of subject 4 in training set.

Fig S5. Relationship between the values of parameter *a* and *k* in Equation 10 in the main text to target’s distance from fixation location.

Fig S6. Locations of the 400 positions where the model can fixate.

Fig S7. Variability of saccadic landing position as a function of saccade target eccentricity.

Fig S8. Fixation location distribution of subjects and models in error trials.

Fig S9. Distribution of saccade amplitude as a function of ordinal position in a sequence of saccades from all correct trials.

Fig S10. Distribution of saccade amplitude as a function of ordinal position in a sequence of saccades from all error trials.

Fig S11. Distribution of fixation duration as a function of ordinal position in a sequence of fixations from all correct trials.

Fig S12. Distribution of fixation duration as a function of ordinal position in a sequence of fixations from all error trials.

Fig S13. Relationship between the change in saccade direction and the preview benefit of the fixation in between.

S1 Text. Calculation of the Posterior Probability Map.

S2 Text. Validation of the ELM Rule.

S3 Text. Visibility Map without Target Location Cue.

S1 Table. Parameters used in the three visual search models.

