## Supporting Information for "Human Visual Search Follows Suboptimal Bayesian Strategy Revealed by a Spatiotemporal Computational Model"

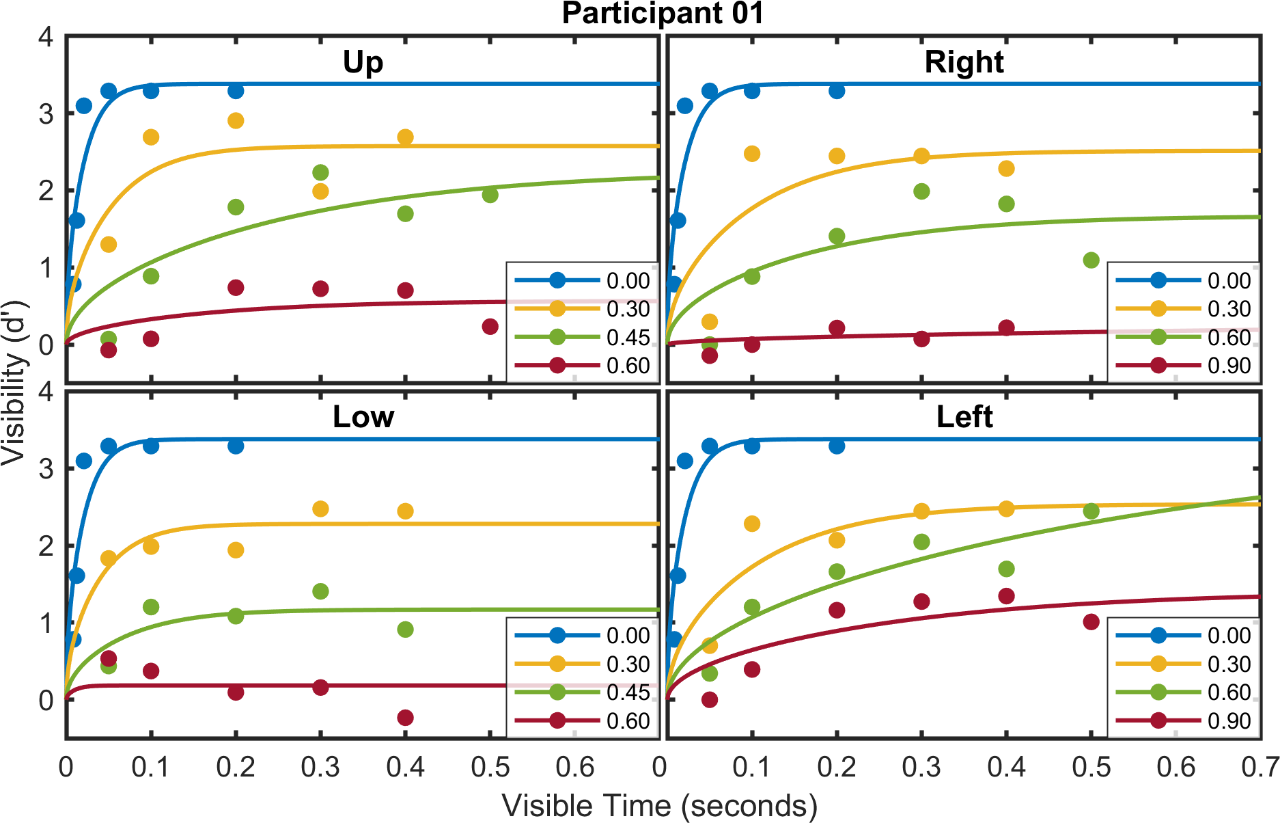

Figure S1. Temporal course of target visibility at four cardinal directions (shown in four subplots) relative to fixation location of subject 1 in training set. Each color (gray scale) represents data measured at different eccentricities (shown in legend in each subplot). The numbers in legend represent the relative distance between measured location to fixation location (the radius of search field is one unit, 0.00 means fixation location). The data measured at fixation location are shown in every subplot. Dots represent raw data and lines are Equation 10 fitted to the raw data measured at each location.

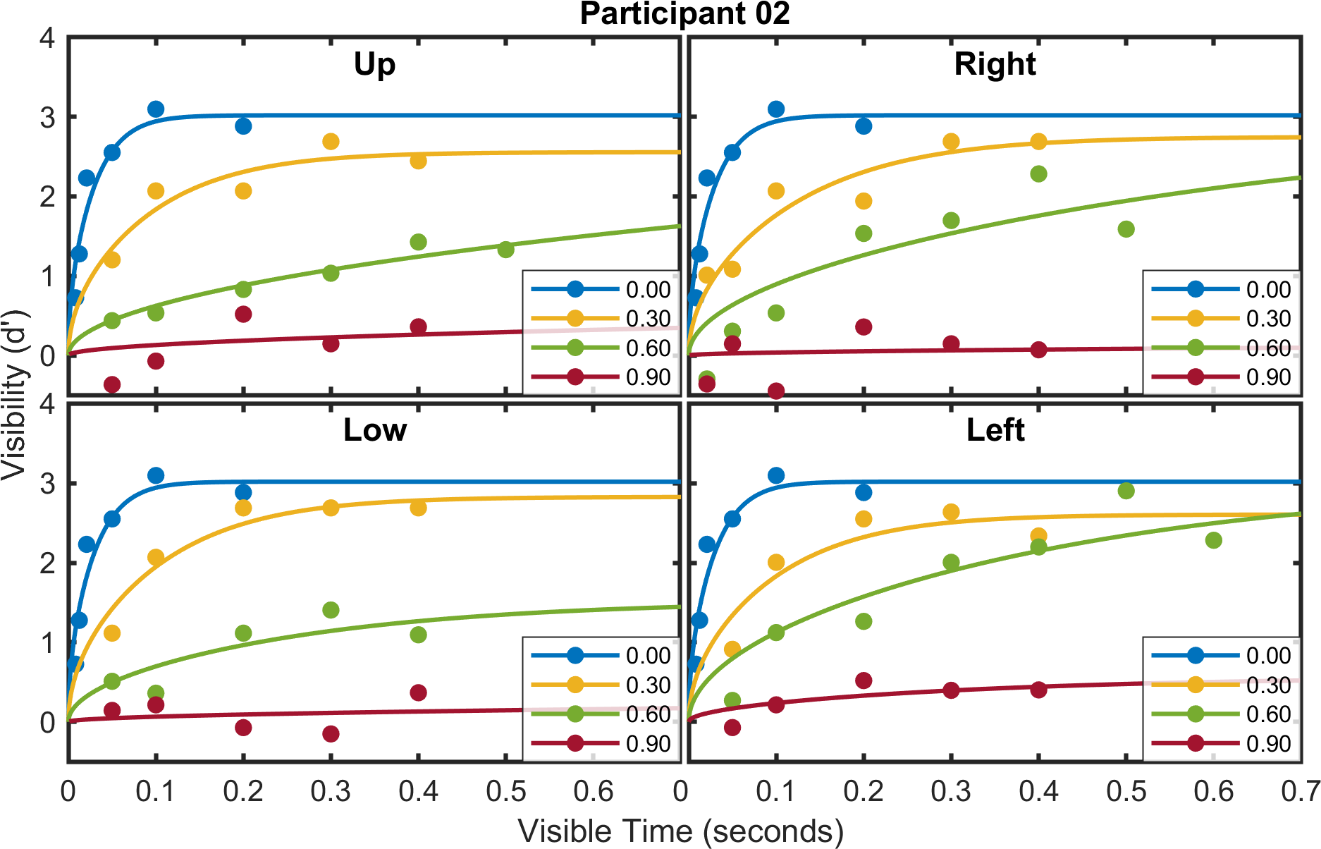

Figure S2. Temporal course of target visibility at four cardinal directions (shown in four subplots) relative to fixation location of subject 2 in training set. Everything else is the same as Figure S1.

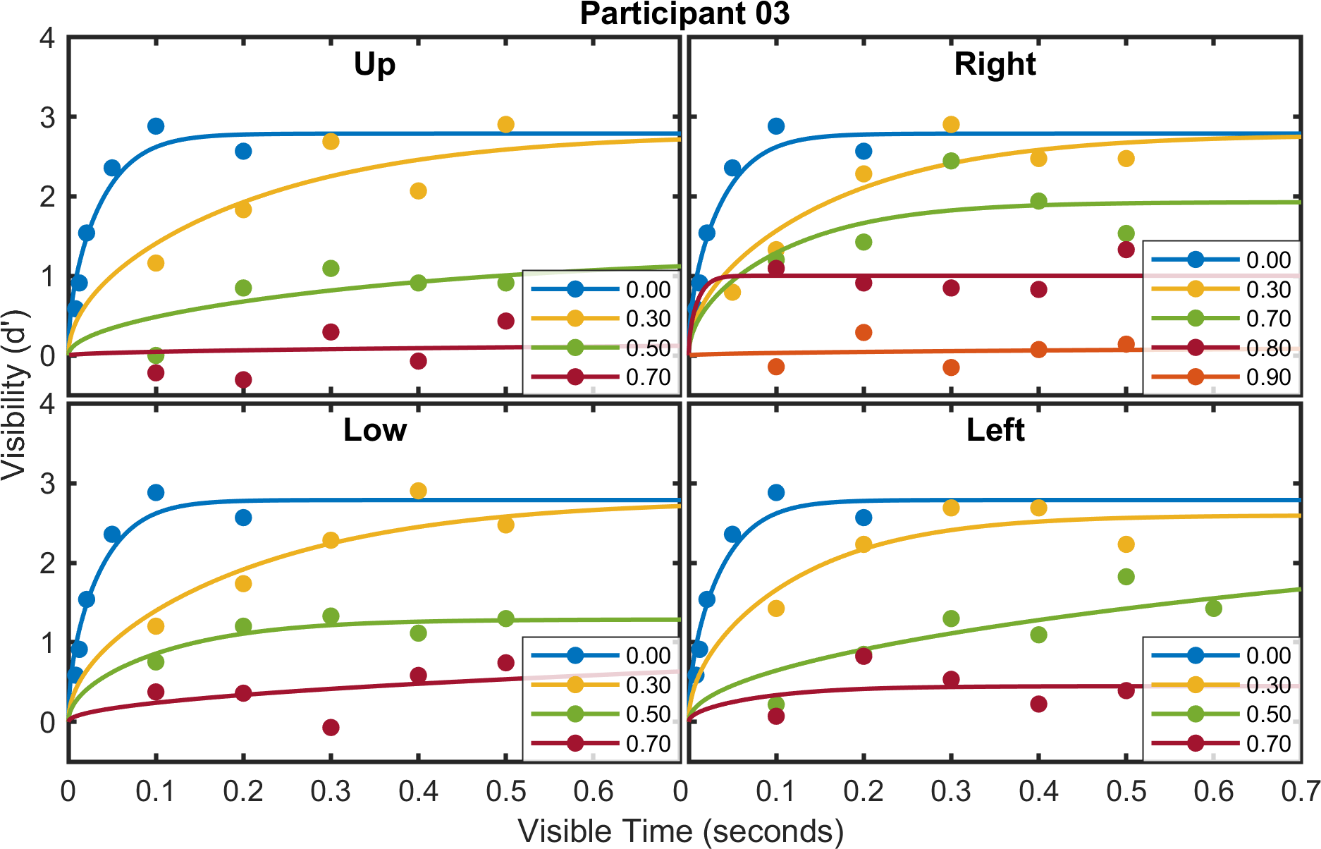

Figure S3. Temporal course of target visibility at four cardinal directions (shown in four subplots) relative to fixation location of subject 3 in training set. Everything else is the same as Figure S1.

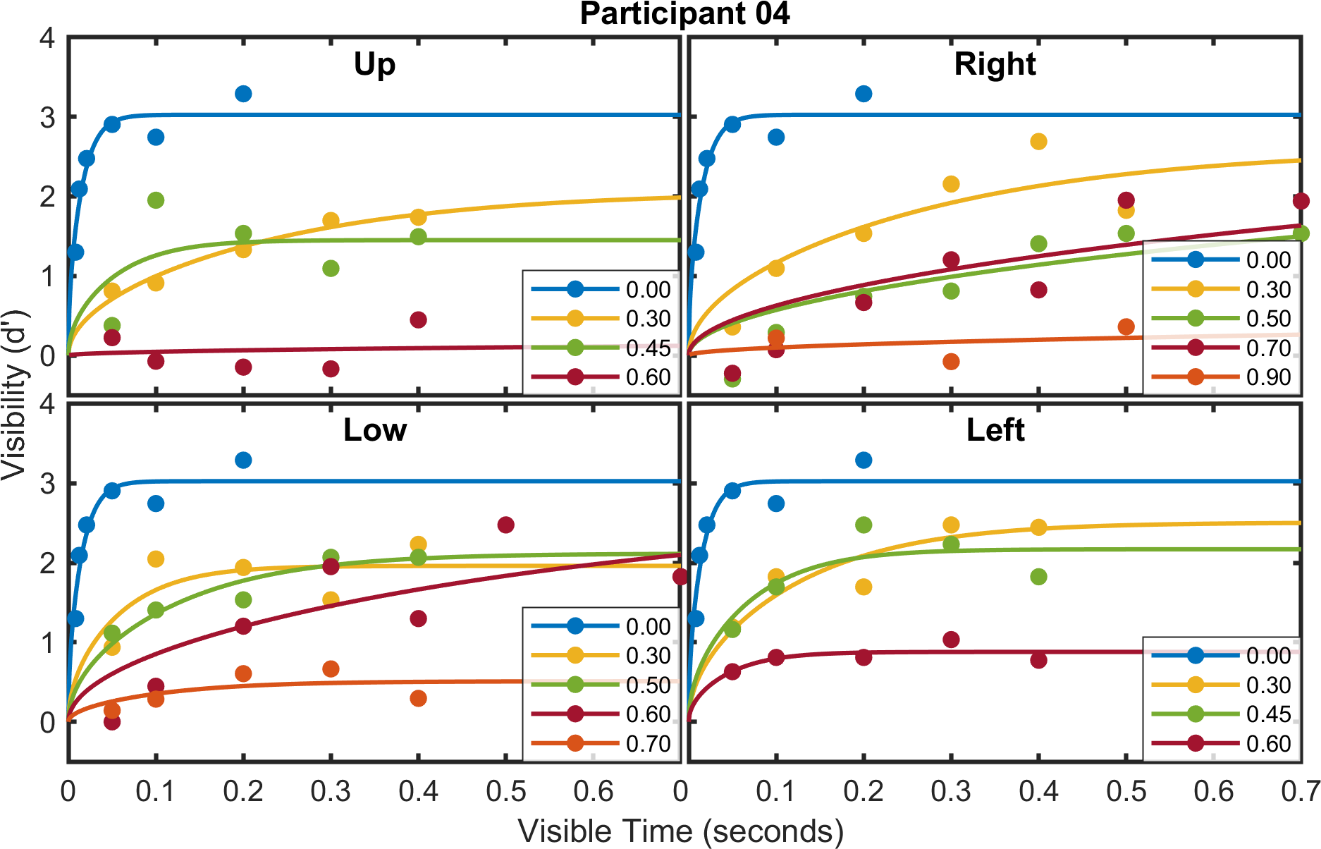

Figure S4. Temporal course of target visibility at four cardinal directions (shown in four subplots) relative to fixation location of subject 4 in training set. Everything else is the same as Figure S1.

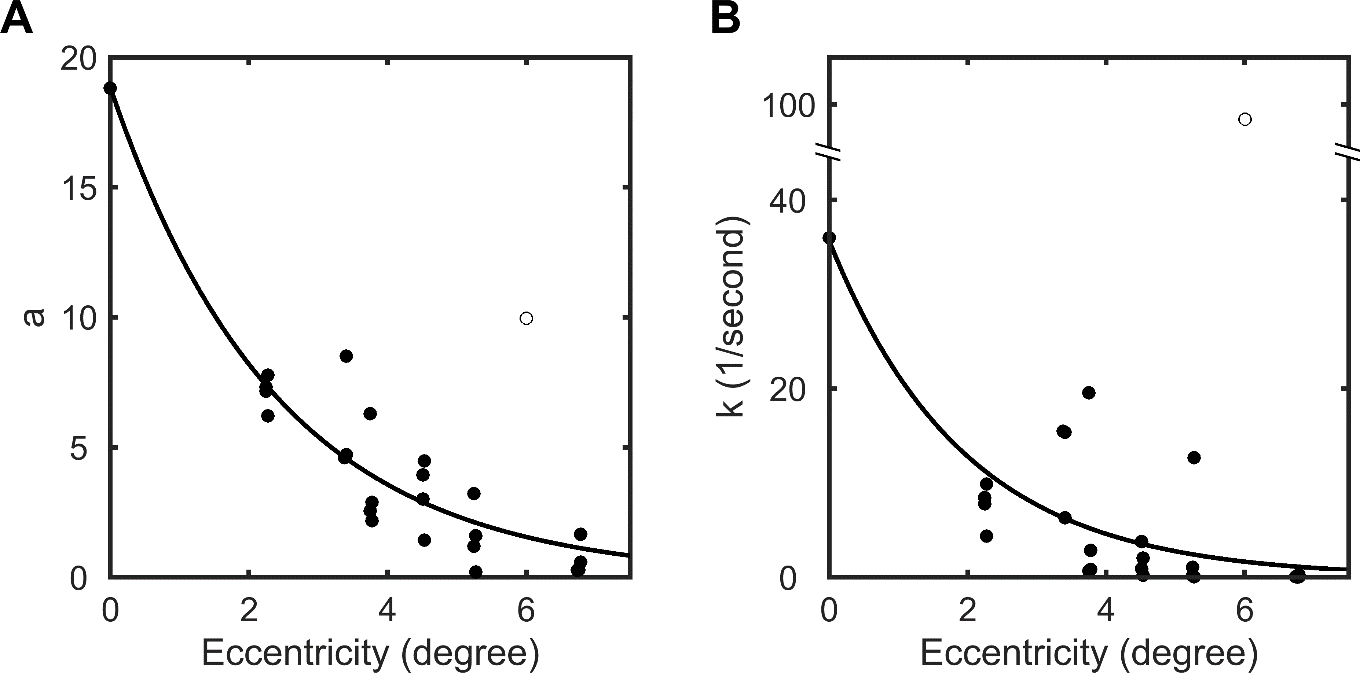

Figure S5. Relationship between the values of parameter *a* and *k* in Equation 10 in the main text to target’s distance from fixation location. Each dot in the plot represents the values of parameter a and k obtained by separately fitting Equation 10 to the time course of target visibility measured at each location. The open circle represents an outlier. The line is an exponential function *y* = *β*_1_ ∙ exp(−*β*_2_ ∙ *x*) fit to the dots (*β*_1_ and *β*_2_ are parameters, the outlier are ignored). Here we grouped target location only by their distance to fixation center and ignored the direction.

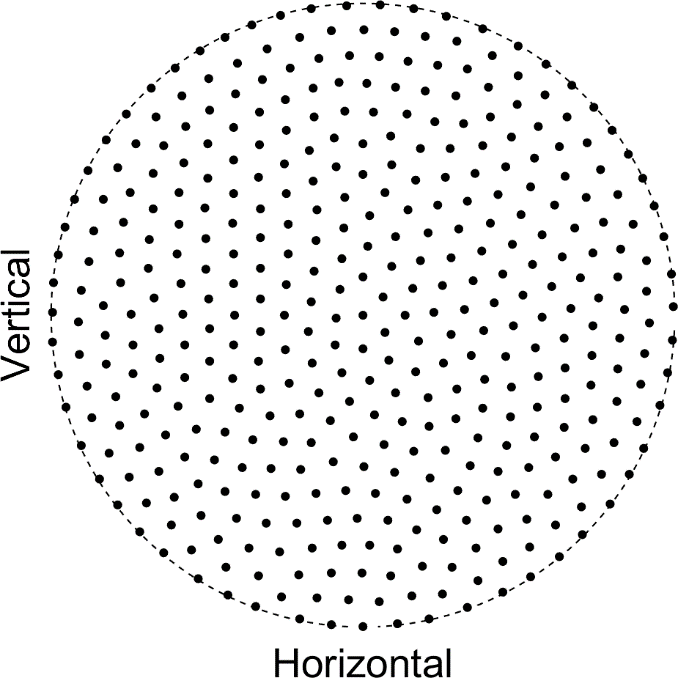

Figure S6. Locations of the 400 positions where the model can fixate. The dashed circle is the boundary of the search area. We generated these positions by simulating 400 dots repelling each other. The repelling force was proportional to the inverse sixth power of the distance between two dots. One dot was fixed at the image center and served as the starting position of the visual search.

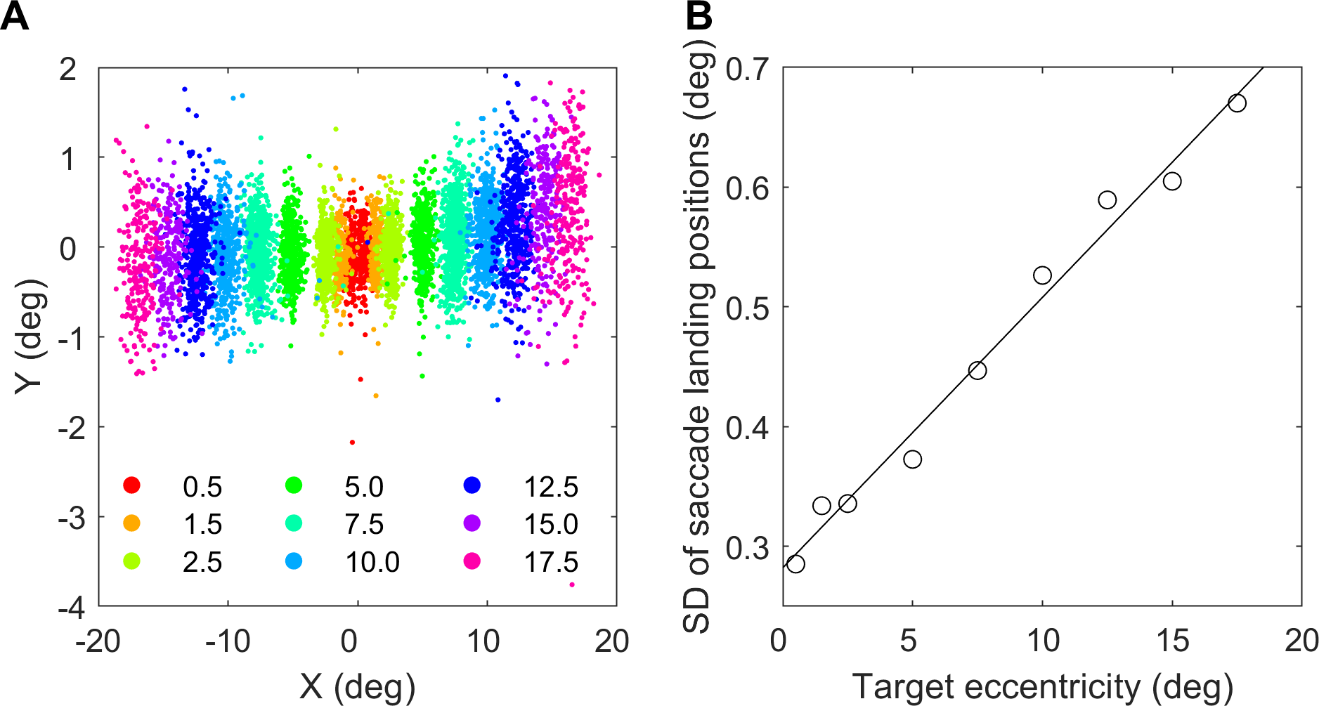

Figure S7. Variability of saccadic landing position as a function of saccade target eccentricity. The two plots are generated by re-analyzing data from Nuthmann et.al, 2016 (Ref [12] in the main text). ***A***. Raw saccade landing position to target at 0.5 – 17.5 degrees (shown in legend) leftward and rightward from initial fixation location. Each dot represents data from one trial. ***B.*** Standard deviation of saccade landing position as a function of target eccentricity. Leftward and rightward of the same target eccentricity are combined. Standard deviation was calculated by Equation 26 in the main text. Circles are experiment data, and the line is the best-fitting line.

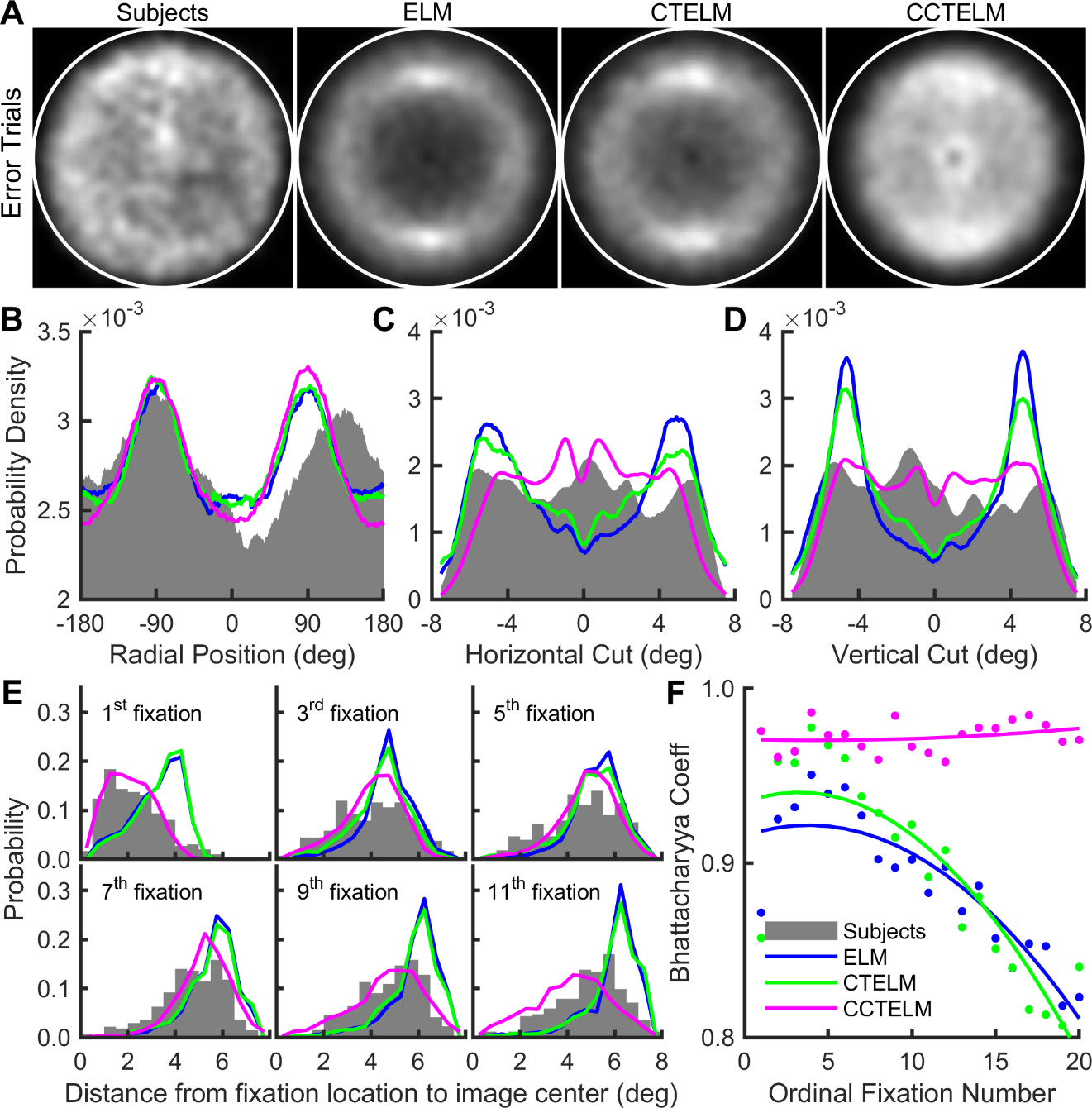

Figure S8. Fixation location distribution of subjects and models in error trials. **A.** Distribution of fixations location in the search field (inside the white circle), lighter means higher density. The densities were obtained by smoothing the scatterplot of the fixation locations by a Gaussian window with a standard deviation of 0.35 degrees (15 pixels) and then normalizing the maximum value in each subplot to one. **B.** Direction histograms of fixation location relative to the image center. The histograms were obtained with a sliding radial window with a width of 45° centered different radial positions (x-axis). Rightward direction is 0° and upward direction is −90°. **C.** Horizontal cuts (through the center) through the fixation densities in panel A. Rightward direction is positive in x-axis. **D.** Vertical cuts (through the center) through fixation densities in panel A. Upward direction is negative in x-axis. In panels BCD area under curve are normalized to one. **E.** Fixation distance distribution to image center selected within the initial 11 fixations after the first saccade. **F.** Bhattacharyya coefficient between models’ and subjects’ distributions of fixation distance to image center of the initial 20 fixations after the first saccade. Dots represent raw data; curves represent quadratic functions fitted to the dots. Legends from panel B to F are shown in bottom-left of panel F.

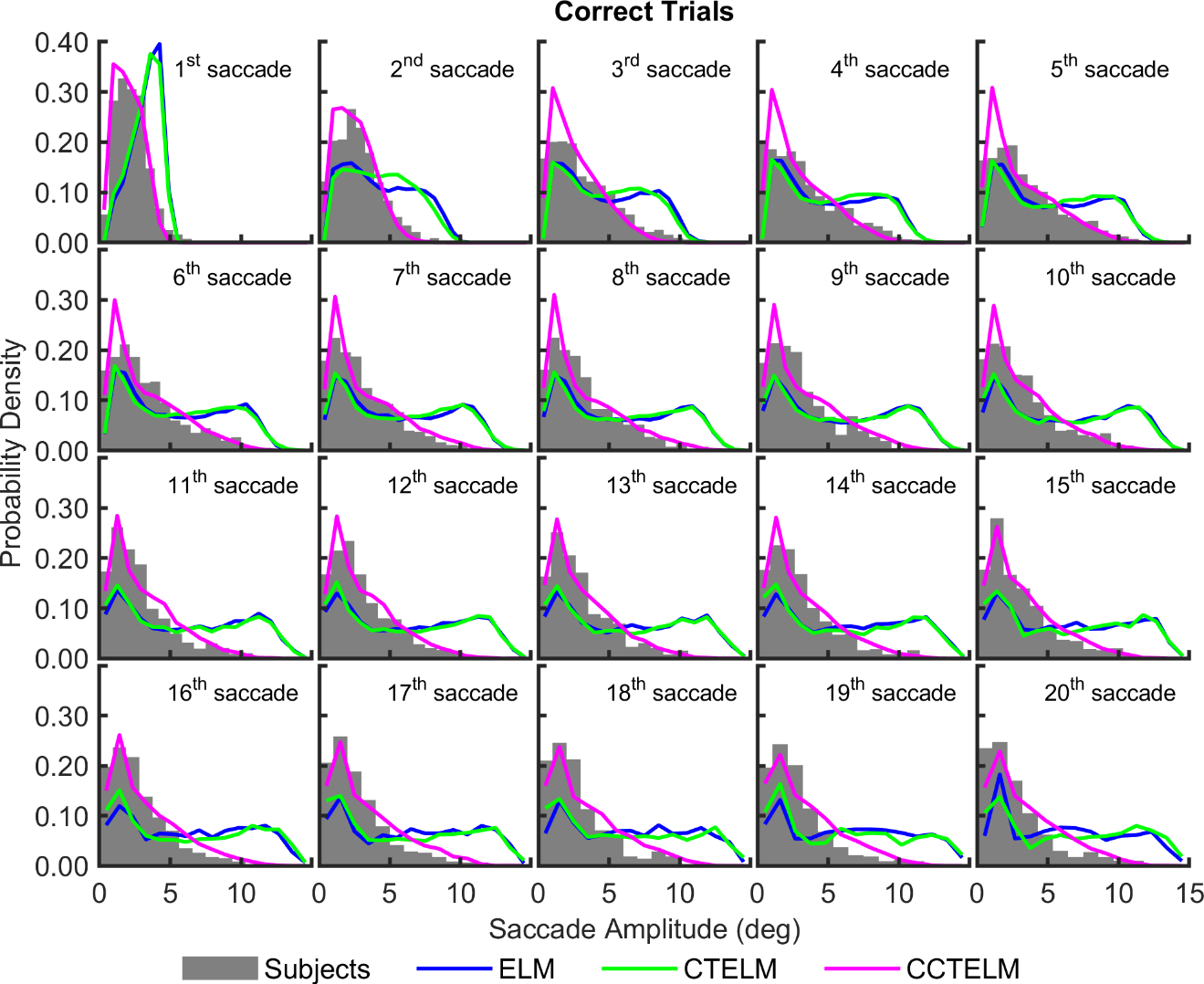

Figure S9. Distribution of saccade amplitude as a function of ordinal position in a sequence of saccades from all correct trials.

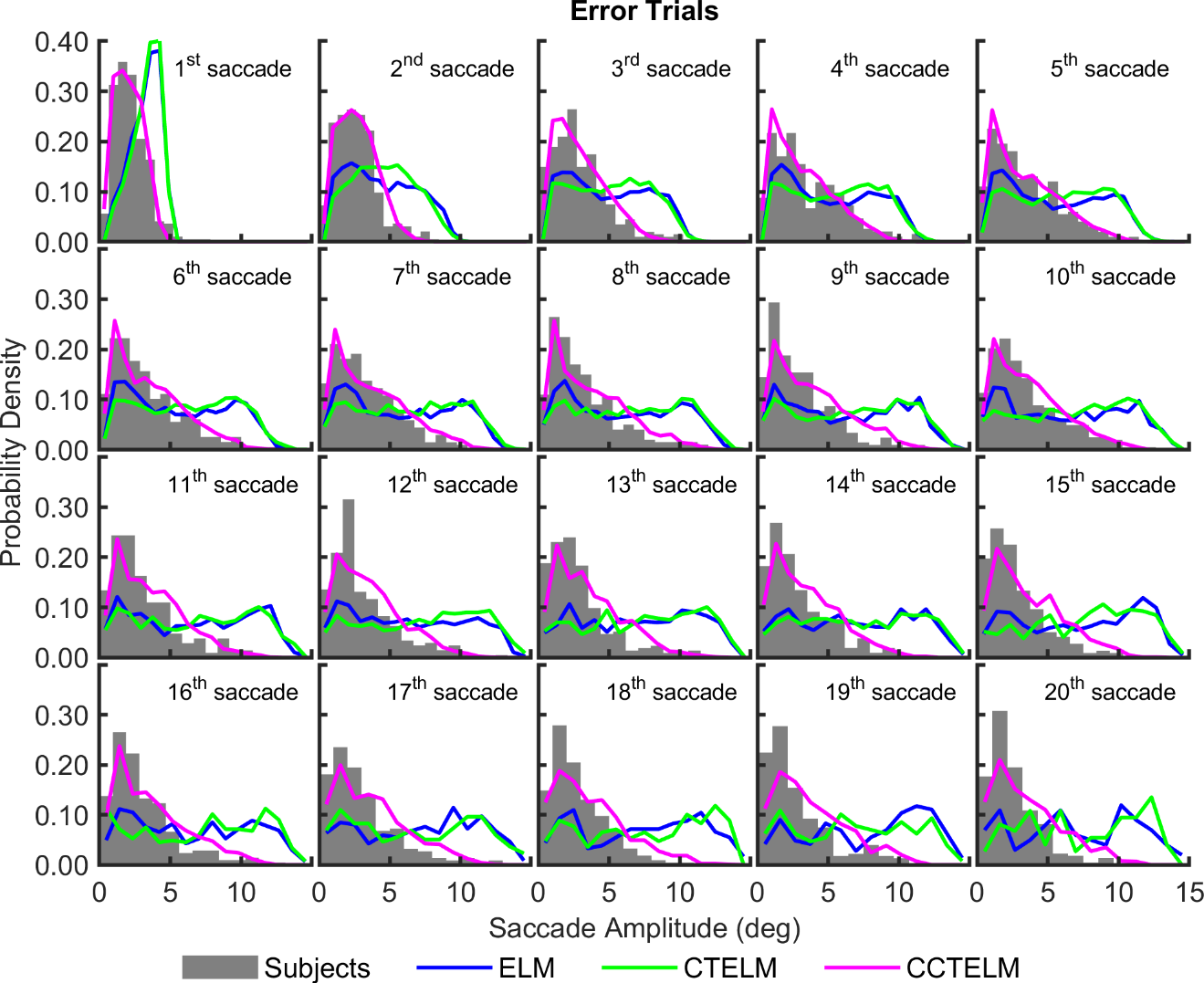

Figure S10. Distribution of saccade amplitude as a function of ordinal position in a sequence of saccades from all error trials.

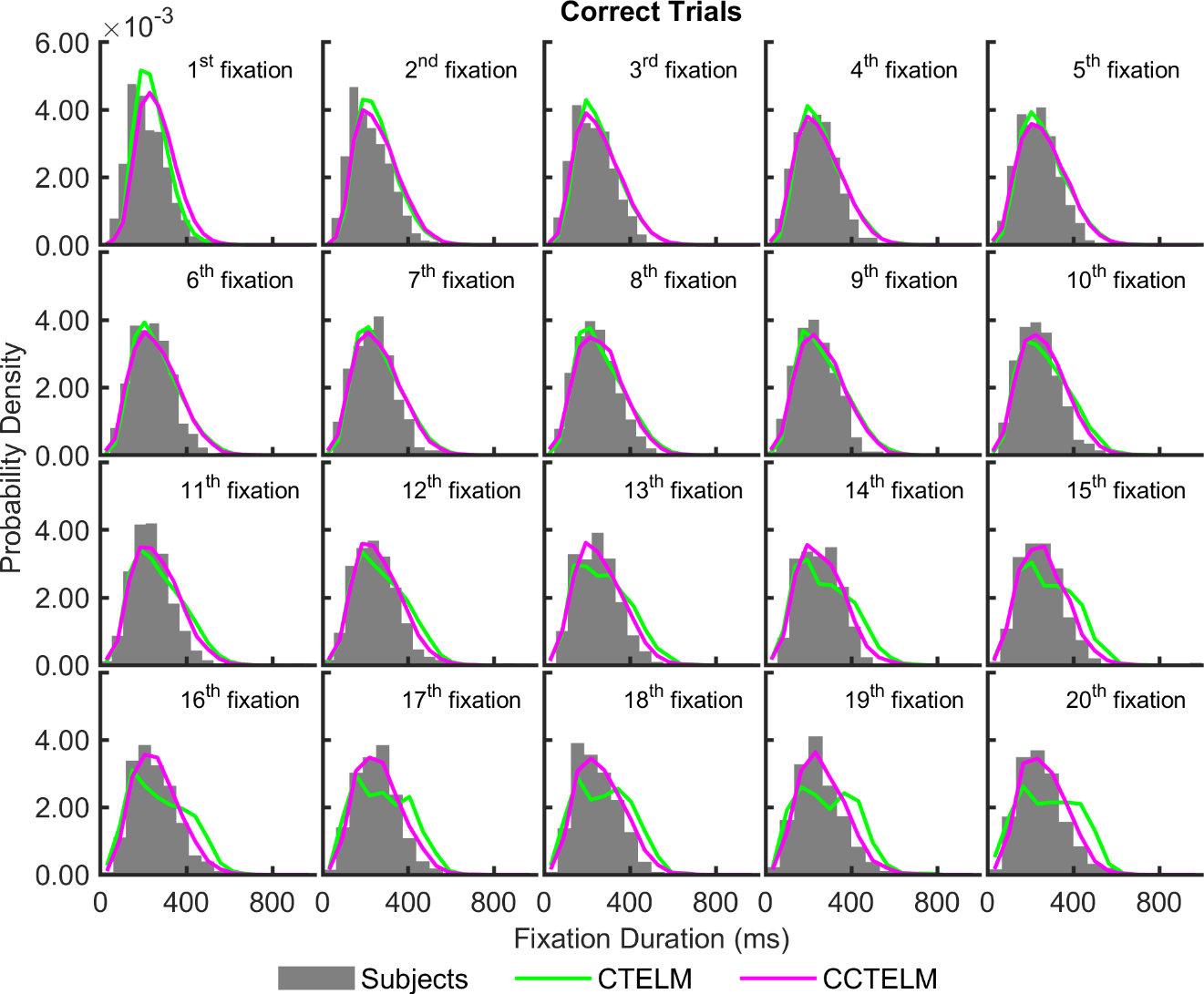

Figure S11. Distribution of fixation duration as a function of ordinal position in a sequence of fixations from all correct trials.

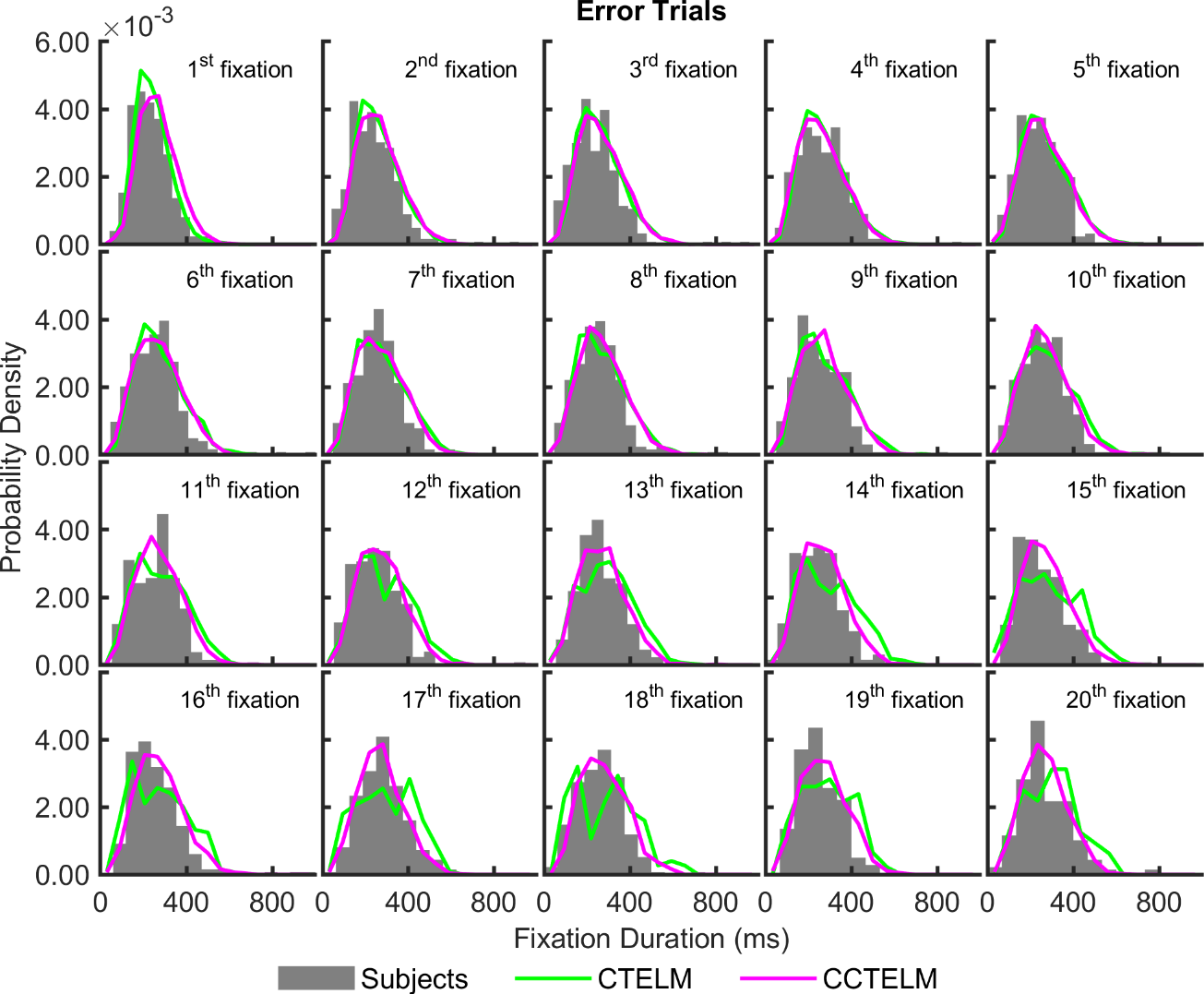

Figure S12. Distribution of fixation duration as a function of ordinal position in a sequence of fixations from all error trials.

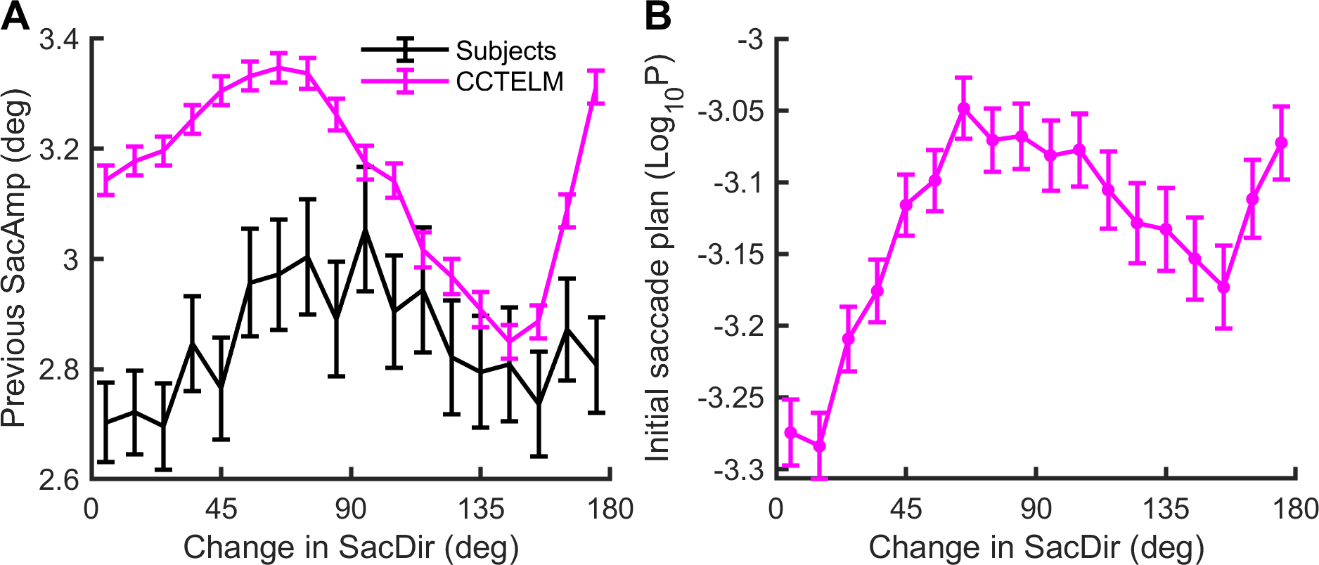

Figure S13. Relationship between the change in saccade direction and the preview benefit of the fixation in between. **A.** Relationship between the change in saccade direction (SacDir) and the first saccade’s amplitude in two consecutive saccades and in all trials. **B.** Relationship between the change in saccade direction and the initial value of the second saccade’s decision process (the posterior probability of target being at current attention location in logarithm scale) in two consecutive saccades and in all trials. Lower initial value was closer to the saccade decision threshold and had larger preview benefit. In A and B error bar shows 95% confidence interval.

**S1 Text**

**Calculation of the Posterior Probability Map**

Here we show the details of derivation of the expression to calculate posterior probability map of target location (Equations 20 and 23 in the main text).

**Model with Unlimited Memory**

According to Equation 19 in the main text, at time *T_F_* after the start of the *F^th^* fixation, the posterior probability of target being at location *i* given accumulated information **W** from all previous fixations is:

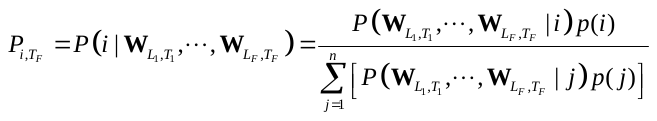

We assume that visual information is independently accumulated at each location during each fixation, so the joint probability in Equation can be simplified to:

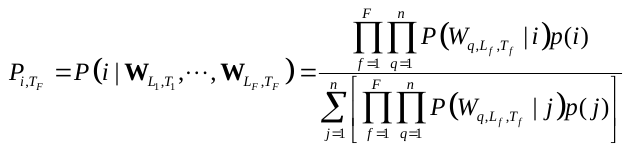

From Equation 7 in the main text we know the accumulated information
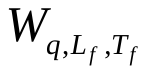
 at location *q* when the *f^th^* fixation is at *L_f_* follows the normal distribution:

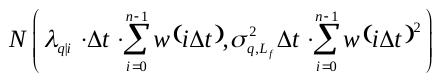

where *λ_q|i_* = 1 if *q* = *i*, otherwise *λ_q|i_* = −1; *T_f_* = *n_t_* ∙ ∆*t*, and
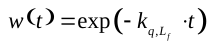
. Note that
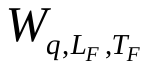
 only contains the accumulated information from the start of the *F^th^* fixation. Information from all previous fixations are stored in
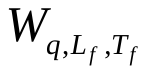
 where *f =* 1⋯*F*−1*,* and they are all used to calculate the posterior probability during the *F^th^* fixation in Equation . Taking the limit ∆*t* →0 and the summation of *w*(*i*∆*t*) becomes integration, and note that
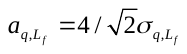
 (Equation 10 in the main text), the distribution becomes:

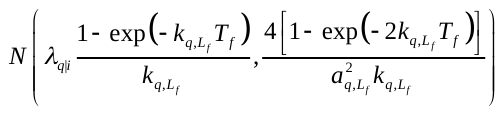

Here the parameters *a* and *k* have subscripts *q* and *L_f_* because their values depend on the relative location between fixation location *L_f_* and the queried location *q* (Equations 12, 13 in the main text). Now we can write the probability density function
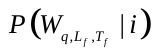
 as:

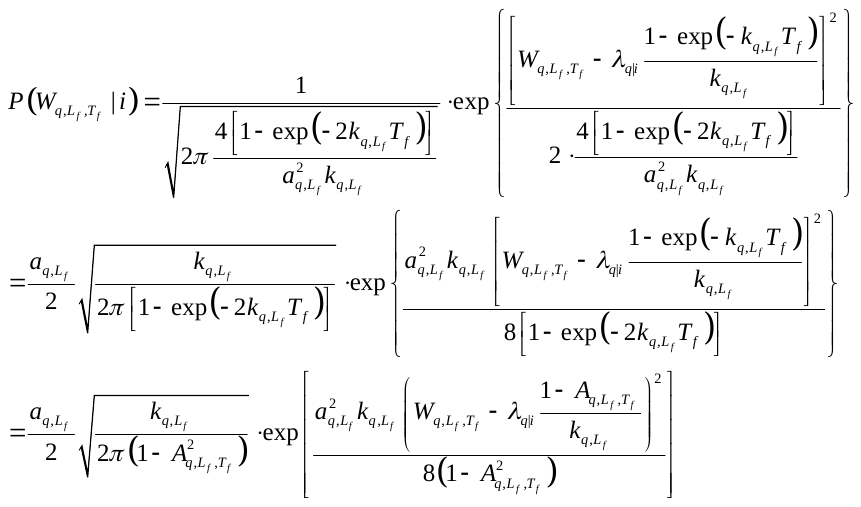

Note in the last step of Equation we define
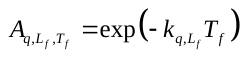
.

Substituting Equation into Equation :

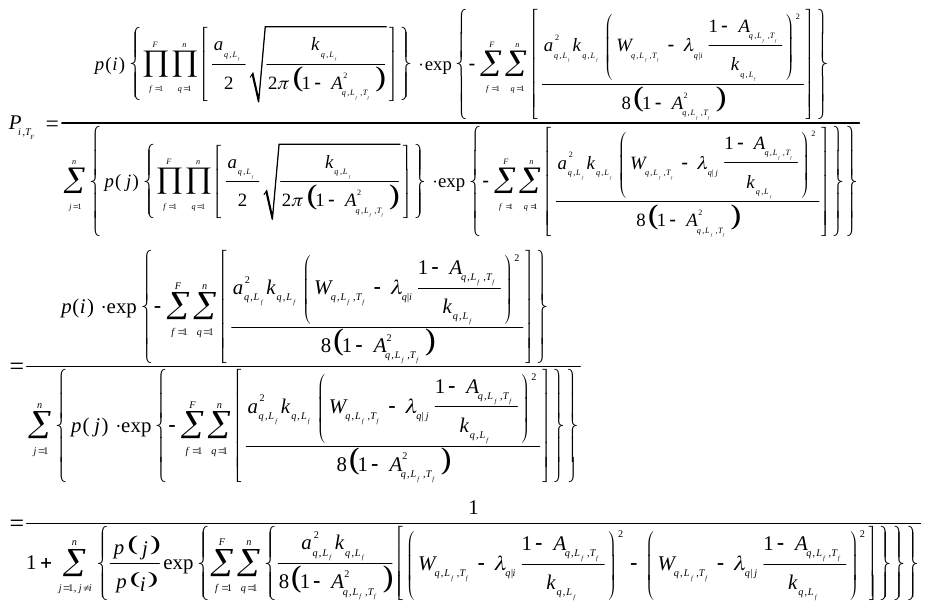

Note that:

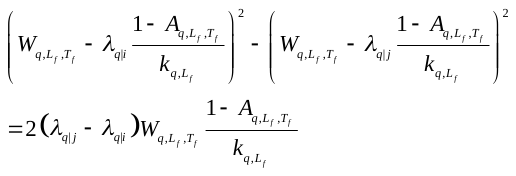

Substituting Equation into Equation , and note that (*λ_q|i_* - *λ_q|j_*) = 0 for *q* ≠ *i* and *q* ≠ *j*, we have:

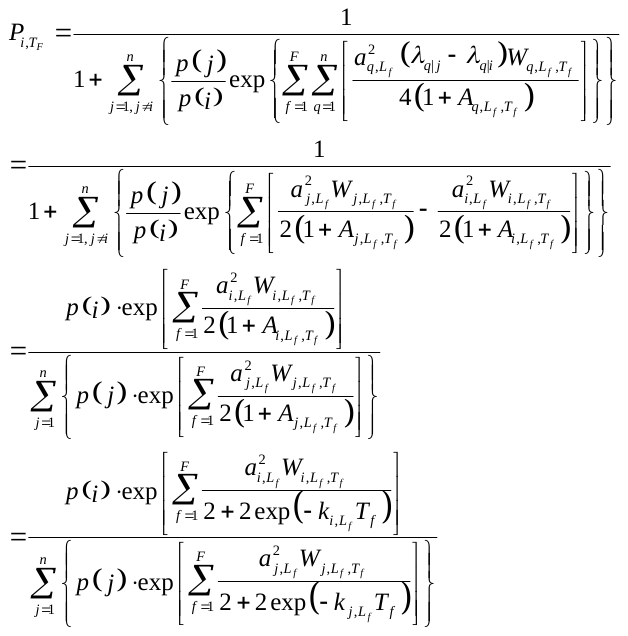

and this is the formula to calculate posterior probability map in Equation 20 in the main text.

**Model with Limited Memory**

Model with a memory capacity of *M* can only integration information from current fixation plus *M*−1 previous fixations, so the earliest fixation that the model could integrate is *f_s_* = max(1, *F*−*M*+1). Therefore, Equation should be written as:

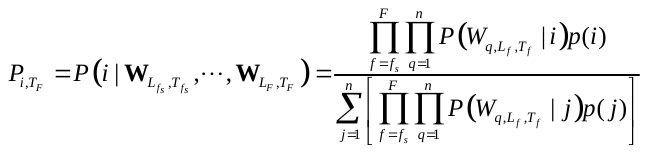

With the same process from Equation to Equation , the posterior probability of target being at location *i* at time *T_F_* after the start of the *F^th^* fixation is:

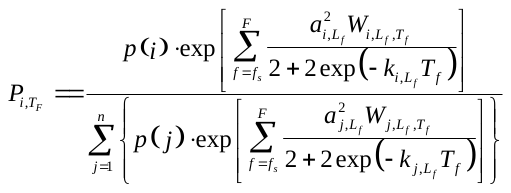

And this is Equation 23 in the main text to calculate posterior probability map for a model with limited memory.

**S2 Text**

**Validation of the ELM Rule**

The calculation of expected information gain of the next fixation in Equation 17 in the main text was derived by Najemnik & Geisler (2009) (Ref [1] in the main txt) without the consideration of time, so the target visibility map was treated as fixed values. However, in our model the target visibility changes according to fixation duration, which is a random variable, so mathematically the derivation no longer holds. Here we used a Monte Carlo simulation to show that Equation 17 still has a high level of accuracy even when the fixation duration varies stochastically.

The Monte Carlo simulation contained 100 sessions. In each session, we generated a random prior probability map of target location and calculated the actual and expected information gain of across all possible fixation locations. At each fixation location, the expected information gain was calculated by Equation 17, and the actual information gain was obtained by repeatedly simulating the fixation for more than 2000 trials until the average information gain changed less than 0.1%. Within each repetition, the target location was randomly chosen according to the prior probability map. We used the same visibility map, saccade threshold parameters, and fixation termination rule as in the CTELM model (see Table S1 for values of the parameters). The duration of the fixations lied in the range of 0.04 to 0.71 seconds, with a mean value of 0.23 seconds.

At the end of each session we could get a map of expected information gain and a map of actual information gain. We then calculated the Pearson correlation coefficient of these two maps. Figure S14 shows the correlation between expected and actual information gain from the results of randomly selected 9 sessions. The mean correlation coefficient across 100 sessions of simulation is 0.923, with a maximum of 0.964 and a minimum of 0.873.

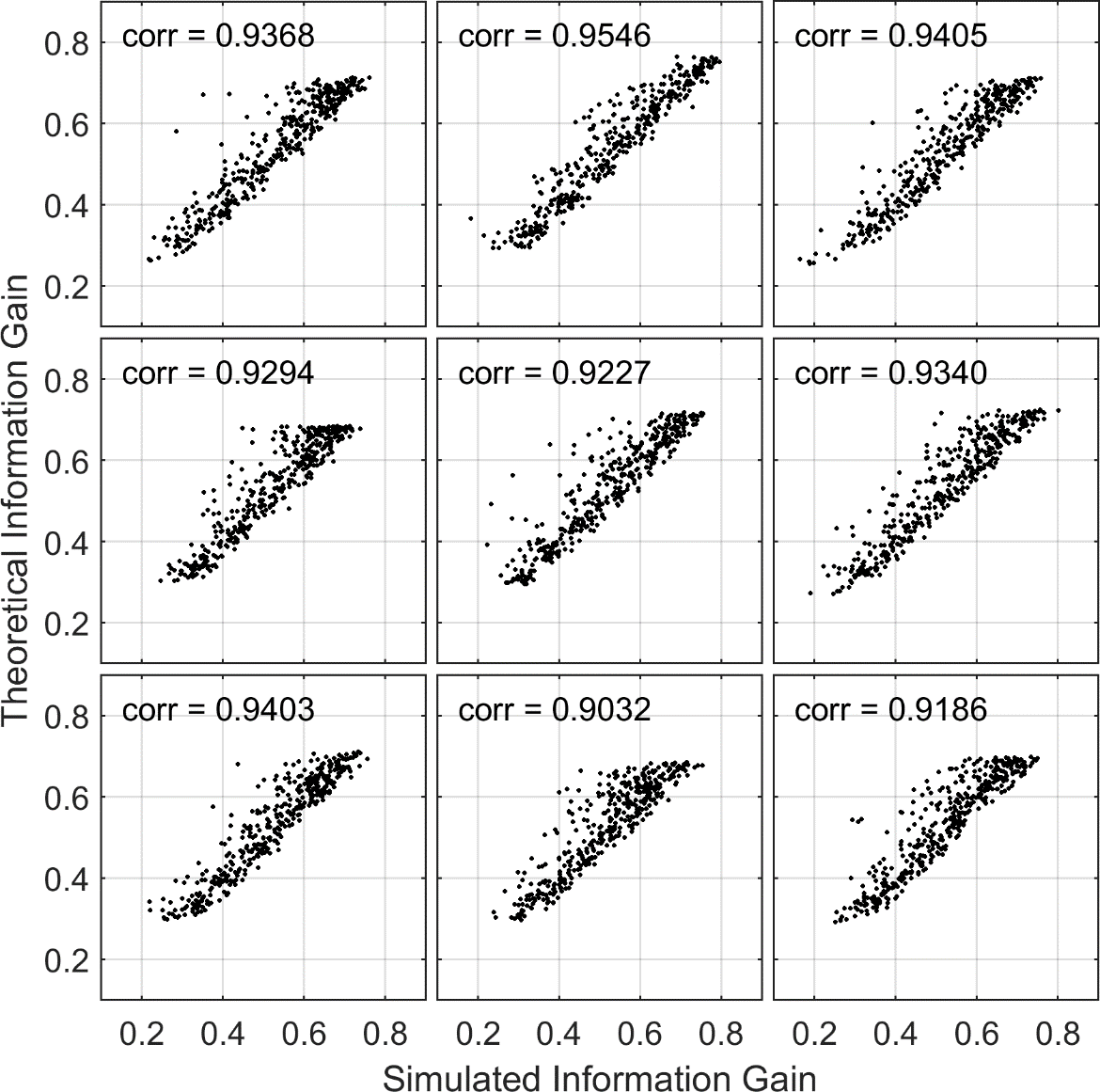

Figure S14. The correlation between expected (vertical axis) and actual (horizontal axis) information gain from the results of randomly selected 9 simulation sessions. Each dot in a plot represents one possible fixation locations, so there are 400 dots in each plot. The Pearson correlation coefficients are shown at the upper left of each plot.

**S3 Text**

**Visibility Map without Target Location Cue**

The visibility map we used in the main text was measured when subjects were cued about target locations. This brings the question of whether this visibility map truly represents the real visibility map subjects used in visual search when there was no clue about target location. We therefore performed the same detection experiment but without the location cue (second version of the detection task in main text) on another four subjects (3 males, age range 19-23 years, all come from the same university as the 4 subjects in the main text, one subject was the author Y.Z.). The target root-mean-squared (RMS) contrast for each subject was determined by the same way as in the main text. These subjects also performed the same visual search task (200-500 trials for each subject, average 340 trials). We identified 21554 fixations and 21824 saccades in total using the same algorithm in the main text. This group of subjects performed close to the train set in the main text, with correct response rate of 89.3%.

The properties of the visibility map were largely the same as visibility map with location cue, except that steady state values were lower (foveal peak visibility = 2.33). At foveal region, visibility peaked within the initial 100 ms, but peripheral regions rise slower and lower peak visibility level (Figure S15).

We used the ELM model (simulated for 10000 trials) to obtain the best possible search performance that this visibility map can offer. The ELM model responded correctly on 88.8% of the trials when the target detection threshold (*θ_T_*) was set to 0.55, but the search performance was much slower. On median, the ELM model needed 17 fixations to correctly find the target, while subjects needed 8 fixations. The overall distribution of fixation number of the ELM model spread more to the slower part than the subjects (Figure S16A). When we split the trials by target eccentricity, the ELM model was slower than subjects under all target eccentricity group in terms of median fixation number (Figure S16B).

Given that the ELM model already searched optimally and did not consider that factors limited search performance, we concluded that the visibility map measured with target location uncued does not allow human level search performance. Therefore, we did not use this visibility map in the main text.

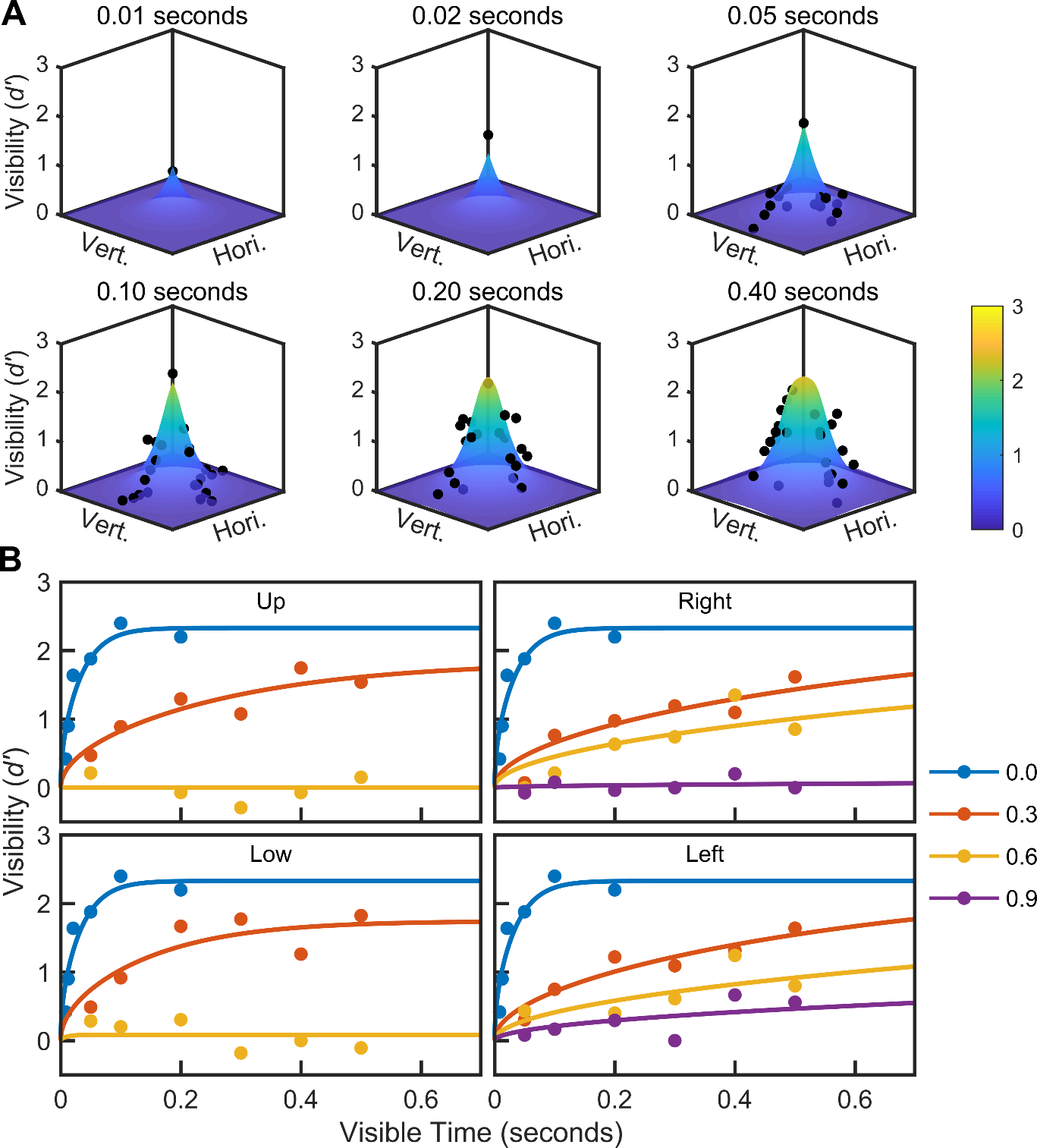

Figure S15. The temporal dynamics of target visibility map when target location is not cued in the detection experiment. Data from all subjects were combined according to measured location. **A.** Each subplot shows target visibility map after different stimulus exposure time in the detection task. Black dots are raw data merged from the four subjects in group one. The surface is the visibility map function (Equations 10, 12,13) fit to the raw data. Not all locations were tested in these exposure time, so the number of black dots may differ in each subplot. Hori/Vert: horizontal/vertical dimension of the search field. **B.** Temporal course of target visibility at four cardinal directions (shown in four subplots) relative to fixation location. Each color represents data measured at different relative eccentricities (shown in legend). Dots are raw data and lines are Equation 10 fitted to the raw data measured at each location. Eccentricity values in legend is normalized with respect to the radius of the search field. The data measured at fixation location (eccentricity = 0.0) are shown in every subplot. For clarity, not all measured locations were shown here.

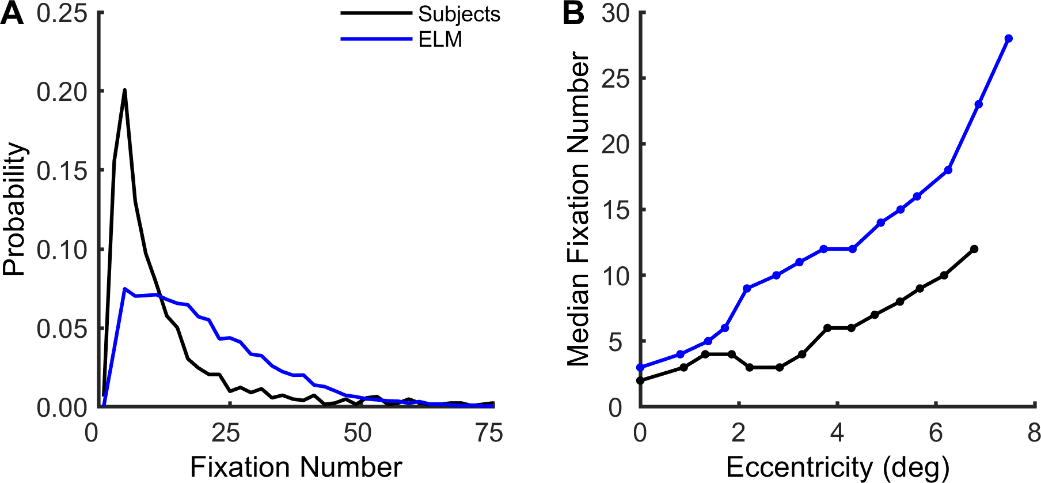

Figure S16. Participants and the ELM model’s search performance in correct trials. The ELM model used the visibility map measured when target location was not cued. **A**. The distribution of the number of fixations needed to find the target. **B**. Median number of fixations as a function of median target eccentricity from image center.

**Parameters used in the three visual search models**

| Model | Parameters | Description | Value | Determination |
| --- | --- | --- | --- | --- |
| Shared | *p*_1_, *p*_2_, *p*_3_, *p*_4_, *p*_5_ | Parameters in visibility map function. | 19.683, 0.00889, 43.195, 0.0153, 1.635 | Fit by GlobalSearch algorithm |
|  | *n* | Number of potential target locations. | 400 | A priori |
|  | *dt** | Time step in seconds. | 0.001 | A priori |
|  | Normal saccade frequency * | The probability that a saccade is triggered by the main decision process. | 97% | A priori, according to low-latency saccade frequency |
|  | Low-latency saccade frequency * | The probability that a saccade is not triggered the main decision process. | 3% | A priori (Munoz et al. 1998) [76] |
|  | Latency distribution of low-latency saccades (seconds) * | The latency from previous saccade decision to the time when eye starts to move in the following saccade. | 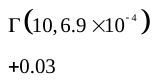 | A priori (Fischer and Ramsperger 1986) [77] |
|  | Eye-brain lag (seconds) * | Time delay from retina to primary visual cortex. | 0.06 | A priori (Nowak and Bullier 1997) [78] |
|  | Saccade lag (seconds) * | Time delay from frontal eye field to eye movement muscle. | 0.03 | A priori (Bruce et al. 2017) [79] |
| ELM | *θ_T_* | Target detection threshold. | 0.240 | A priori (Fit by human correct response rate) |
| CTELM | *q*_1_, *q*_2_, *q*_3_ | Parameters in saccade threshold function. | -9.972, 0.485, 1.020 | Fit by genetic algorithm |
|  | *θ_T_* | Target detection threshold. | 0.982 | A priori (Fit by human correct response rate) |
| CCTELM | *q*_1_, *q*_2_, *q*_3_ | Parameters in saccade threshold function. | -12.864, 0.367, 0.8164 | Fit by genetic algorithm |
|  | *c* | Parameters in saccade amplitude penalty function. | 386.34 | Fit by genetic algorithm |
|  | *θ_T_* | Target detection threshold. | 0.952 | A priori (Fit by human correct response rate) |
|  | *M* | Number of previous fixations that the model can keep in memory. | 8 | A priori (Fit by sequential eye movement metrics) |
|  | *m*, *b* | Linear regression coefficients between standard deviation of saccade landing position and saccade target eccentricity. | 0.0452, 0.364 | A priori, obtained from data in (Nuthmann et al. 2016) [12] |

Table S1*.* Parameters used in the three visual search models in the main text. The value of a priori parameters were determined prior to fitting of other parameters by genetic algorithm. Shared parameters were used in all models except specifically indicated ones. The number after a reference is the reference number in main text.

* Not used in the ELM model.
